# Chromosomal plasticity can drive rapid adaptation in bacteria

**DOI:** 10.1101/2024.12.11.627208

**Authors:** Luis Vega-Cabrera, Ole Skovgaard, Macarena Toll-Riera

**Author notes:** Institut de Biologia Evolutiva (CSIC-UPF), Barcelona, Spain.

## Abstract

Understanding the molecular mechanisms underlying rapid adaptation to stress is a fundamental question in evolutionary biology. A common mechanism is copy number changes, which can affect single genes, larger genomic regions, or even entire chromosomes in the case of eukaryotes (aneuploidy). Bacterial species with genomes comprising more than one chromosome are common, yet the role of adaptive copy number changes affecting entire chromosomes is unknown. Here, by using a bacterium with two chromosomes (chrI and chrII) as model system and combining evolution experiments in the presence of severe nutrient stress with whole-genome sequencing, we show that chromosomal copy number changes can drive rapid adaptation. We detected increased copy number of the entire chrII in almost half of the sequenced clones, isolated from twelve independent populations. In most cases, copy number increases were driven by mutations in the replication protein of chrII. A higher chrII copy number is likely to be beneficial because chrII codes for the operon responsible for metabolizing D-gluconic acid, the only carbon source present in the medium. Our results revealed a novel mechanism for rapid adaptation in bacteria. Given the prevalence of bacteria with secondary chromosomes, and the enrichment of secondary chromosomes in genes associated with interaction with the environment, this may be an important mechanism for adapting to novel environments that has been overlooked.

## INTRODUCTION

Organisms live in environments that are constantly changing. To survive, organisms must be able to respond rapidly to these changes. This can be achieved, for example through physiological responses. However, if the stress persists over time, more durable genetic mechanisms are required. A common genetic mechanism for rapid adaptation to severe stress is copy number changes, which includes copy number variations (CNVs) and aneuploidies. CNVs involve the amplification or deletion of genes or larger segments of DNA, while aneuploidies affect entire chromosomes^1,2^.

CNVs usually change the expression of one or a few genes^3^. Increased expression can be selected for under conditions in which gene dosage is beneficial for growth^4–11^. Examples of adaptive CNVs have been found throughout the tree of life. CNVs affecting the membrane transporters HXT6/7 and GAP1 are selected in yeast facing glucose and nitrogen limitation respectively^9,12,13^ and oncogene amplification is common in cancers because it promotes proliferation^14^. In bacteria, duplication of one or multiple genes occurs at a much higher rate than point mutations^15,16^, and drives adaptation to nutrient limitation^17,18^, to novel environments^19^, and the evasion of the host immune response^20^.

In contrast to CNVs, aneuploidy alters the expression of thousands of genes^3^. Although aneuploidy is a priori detrimental^21–25^, it is widespread in nature and has been found in yeast, in mammals and in plants^23^. Experimental evolution studies have shown that aneuploidy is a common mechanism in microbial eukaryotes to rapidly adapt to stressful conditions^9,13,26–29^. For instance, yeast exposed to starvation stress gained an additional copy of the chromosome that encodes the transporter responsible for the uptake of the scarce nutrient^9,13^.

Aneuploidy and CNVs are simple and quick solutions to abrupt and severe stress^29^, but because they are costly^3,23,30^, they are typically lost once a less costly and more stable solution emerges, such as a point mutation or the duplication of a single gene^1,2,29^. For example, the aneuploidies gained by *Saccharomyces cerevisiae* in response to heat stress were lost when the stress disappeared, but replaced by less costly solutions if the stress persisted^29^. Similarly, the multiple copies of a CNV acquired when *Candida albicans* was exposed to antifungals rapidly reverted to the ancestral copy number when the antifungals were removed^10^. Because of their usually short life, the role of aneuploidies and CNVs in adaptation may be underestimated^26^.

About one-tenth of the bacterial species have their genomes segmented into a main chromosome and at least one additional replicon larger than 350 kb^31^. Additional replicons encoding essential genes are classified as chromids or secondary chromosomes, and their replication and partitioning systems have a plasmidic origin^31,32^. The advantages of multipartite genomes are not fully understood. However, secondary chromosomes appear to acquire horizontally transferred genes at a high rate and are enriched in genes involved in interacting with the environment, which may be crucial for adapting to novel or changing environments^33^. We hypothesize that the partitioning of the genome into multiple replicons may facilitate the variation of the copy number of secondary chromosomes in response to stress.

Here we used a marine bacterium with a multipartite genome, *Pseudoalteromonas haloplanktis* TAC125, to investigate whether copy number changes in a secondary chromosome play a role in bacterial adaptation to stress. *P. haloplanktis* TAC125 is a fast-growing cold-adapted bacterium with two chromosomes (chrI: 2942 genes, chrII: 546 genes) and two plasmids^34,35^. chrII is circular, replicates unidirectionally, and was probably already present in the last common ancestor of the *Pseudoalteromonas* genus^36,37^. Approximately 20% of the genes encoded in chrII resemble plasmid-encoded genes, and it maintains a plasmid-like replication system, suggesting that it originated from a plasmid that acquired essential genes^34^. In a previous study, we adapted *P. haloplanktis* to nutrient stress (low concentrations of D-gluconic acid) and serendipitously found a clone that carried an extra copy of chrII but a single copy of chrI^38^. In this study, we systematically investigated whether increases in the copy number of chrII are a common mechanism to adapt to severe nutrient limitation. We found that almost half of the sequenced clones had an increased copy number of the entire chrII, and six additional clones had large CNVs affecting the gluconate operon located on chrII. D-gluconic acid is the sole carbon source in our experiment, likely explaining the beneficial effect of the changes in copy number we observed in chrII. Our findings show that increases in the copy number of a secondary chromosome can mediate rapid adaptation to stress in bacteria.

## RESULTS

### Rapid adaptation to nutrient stress

*P. haloplanktis* TAC125 has a multipartite genome with a main chromosome and a unidirectionally replicating secondary chromosome, chrII^34^. Replication of chrII is controlled by a repA-like protein^34^. In a previous study, we discovered that the number of complete copies of chrII can increase^38^. Specifically, we adapted 12 replicate populations of the wild-type strain to low nutrient (D-gluconic acid) concentration for 200 generations (Supplementary Fig. 1). Whole genome sequencing of the clone that had a better growth relative to the wild-type strain revealed that it had an additional complete copy of chrII, whereas it had only a single copy of chrI^38^.

To investigate whether variations in chromosome copy number could drive rapid adaptation in bacteria, we revisited our previous evolution experiment and systematically studied all endpoint populations^38^. We isolated ten clones from each of the twelve endpoint populations and we compared their growth with that of the wild-type clone used to start the evolution experiment. Fitness increased despite the short duration of the experiment. All adapted clones reached a higher population density, had a higher growth rate, or showed a shorter lag phase than the wild-type clone (Supplementary Fig. 2). The adapted clones exhibited distinctive growth curves, which we classified into monophasic and biphasic growth curves. A total of 78 clones exhibited a monophasic growth curve, the typical bacterial growth curve. Conversely, 42 clones had a biphasic growth curve, which consisted of two consecutive growth phases, similar to the growth of the wild-type clone (Supplementary Fig. 2). Clones harboring one or the other type of growth curve coexisted in almost all populations (Supplementary Fig. 2, Supplementary Table 1, Data file 1). The presence of two distinctive growth patterns associated with higher fitness suggests that there are at least two mutational routes of adaptation to low nutrient concentration.

### An increase in the copy number of the entire chromosome II is a frequent adaptive mechanism to nutrient stress

To identify the genomic basis of adaptation to low nutrient concentration we sequenced clones isolated from endpoint populations. Since the growth curves suggested the existence of two mutational routes, for each of the 12 endpoint populations we randomly selected four or five clones based on the proportion of monophasic/biphasic growth curves found in the specific endpoint population (Supplementary Fig. 2, Supplementary Table 1). We sequenced the genome of 56 clones in total.

We found that over half of the sequenced clones (54%, 30/56 clones) had proportionally more reads overlapping chrII than chrI (Fig. 1A, Supplementary Figs. 3 and 4). Of those clones with higher coverage in chrII than in chrI, 83% (25/30 clones) had an increased read depth that covered the entire chrII, indicating the presence of more than one copy of chrII (↑chrII). In the case of the remaining five clones (1B, 1C, 1D, 3D, 6D), the increase in read depth did not cover the entire length of chrII, suggesting the presence of a CNV that amplifies a segment of chrII between positions 132,527 and 619,758 (CNV_L_-chrII) (Fig. 1A and 1B). We used two different approaches to confirm our observations. First, we used quantitative PCR (qPCR) performed on genomic DNA to determine the copy number of two genes located at distant locations in chrII: *parA*, falling outside the region that showed a partial increase in the copy number (CNV_L_-chrII), and *gluc*, which falls within the aforementioned region (Supplementary Fig. 5). Second, we used marker frequency analysis (MFA)^39,40^ on four selected clones (Fig. 1C). Both qPCR and MFA confirmed the increases in the copy number of the entire chrII (↑chrII) and CNV_L_-chrII. For most clones with copy number changes in chrII (↑chrII and CNV_L_-chrII) the copy number is between 1.25 and 2 (Fig. 1A, Supplementary Figs. 4 and 5).

**Figure 1.**
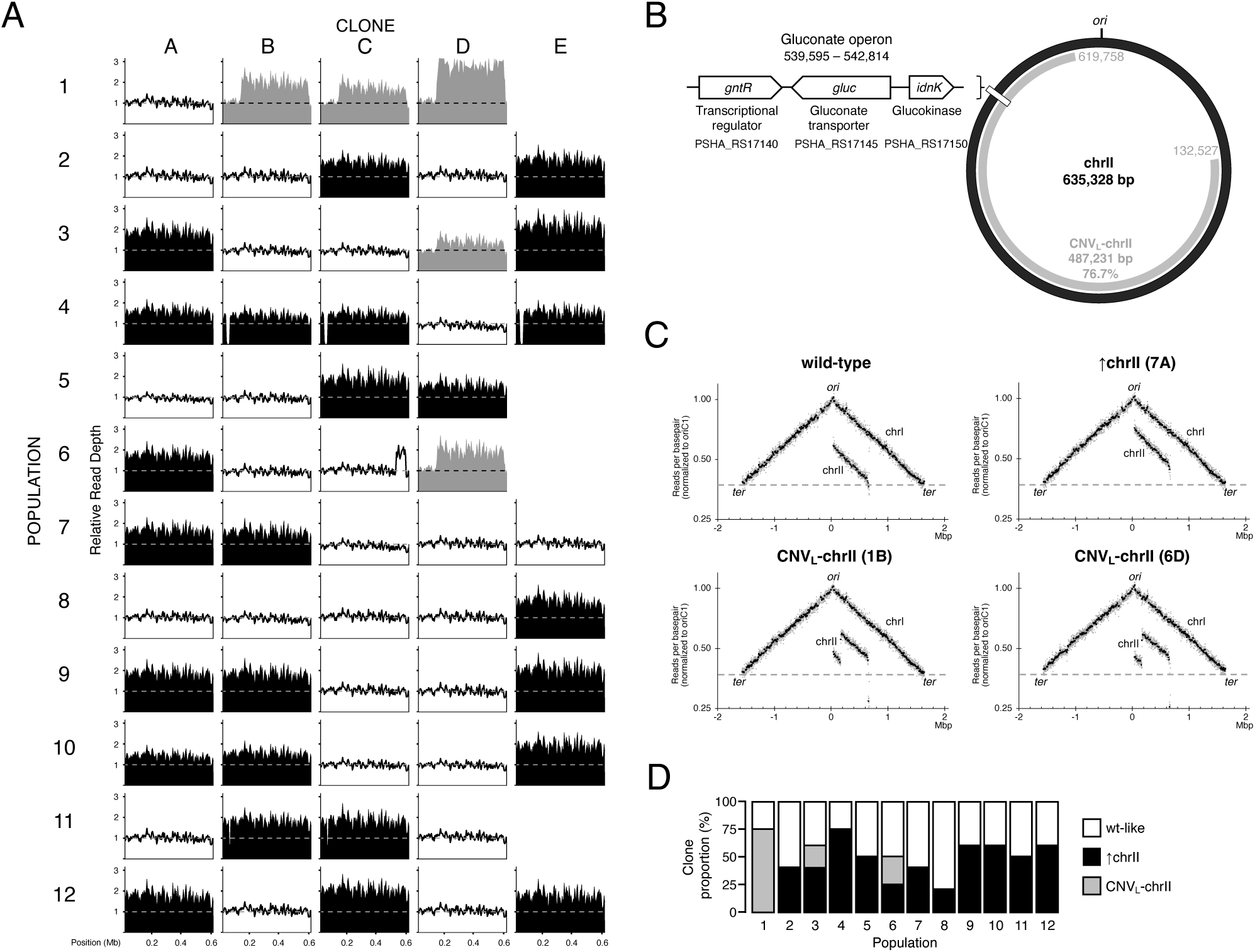
Copy number changes affecting chrII across sequenced clones. A: chrII mean read coverage normalized to chrI for all sequenced clones. Read depth calculated for 5 kb windows. A dashed line indicates the expected relative coverage when no copy number increments are present. White indicates wild-type like clones, black indicates clones that have an increased copy number of the entire chrII and grey depicts clones with CNV_L_-chrII. B: Schematic representation of chrII (black) and the genomic region affected by CNV_L_-chrII (gray). A white rectangle marks the region 539,595:542,814 that contains the operon [*gntR*, *gluc*, *idnK*] to metabolize D-gluconic acid. C: Marker Frequency Analysis (MFA) in exponential phase of the *P. haloplanktis* TAC125 wild-type clone and representative evolved clones. The number of reads (normalized against stationary-phase) is plotted against the chromosomal location. Mean read values of 10,000 bp (black) and 1,000 bp (gray) windows are shown relative to the origin of replication in chrI (oriC1). D: Proportion of sequenced clones from each population that have an increase in the copy number of chrII and CNV_L_-chrII.

We detected increases in the copy number of the entire chrII (↑chrII) in all 12 populations, meaning that this mutation emerged independently and repeatedly in all populations (Fig. 1D). This highly parallel evolution suggests that the increase in chrII copy number is adaptive. chrII encodes an operon responsible for the transport and metabolism of D-gluconic acid (or gluconate), which is composed of three genes: a putative transcriptional repressor for gluconate utilization (PSHA_RS17140), a putative gluconate transporter (PSHA_RS17145), and a putative gluconokinase (PSHA_RS17150) (Fig. 1B). The operon is located between positions 539,655 and 542,184, a genomic region that overlaps with the increase in read depth both in clones with increased copy number of the entire chrII (↑chrII) and CNV_L_-chrII (Fig. 1B). Amplification of the gluconate operon is probably beneficial for growth in limited concentrations of D-gluconic acid as the only available carbon source^38^.

Taken together, our results suggest that copy number changes in chrII (↑chrII and CNV_L_-chrII) are a common strategy to rapidly adapt to nutrient stress by putatively facilitating gluconate uptake.

### The molecular mechanisms underlying the increase in copy number of the entire chrII

We then sought to identify the molecular mechanism responsible for the amplification of the entire chrII (↑chrII). We identified 77 mutations among the 56 clones sequenced. Of these, 58% are in chrI, 42% in chrII and we found no mutations in either of the two plasmids (Supplementary Table 2, Data file 1). The most frequent mutations are point mutations (33) and indels (28), although large deletions and duplications are also common (16) (Supplementary Table 2, Data file 1). All point mutations except one are missense variants. Only one mutation falls in an intergenic region, between the genes PSHA_RS04970 and PSHA_RS04975.

All sequenced clones had at least one mutation, with an average of 1.375 mutations per clone. There are nine genes hit by mutations and three of them, *repA*, *tolC and TBDR*, are repeatedly mutated in several populations (Fig. 2). This suggests that mutations in these three genes are adaptive. The most mutated gene in our experiment is *repA*, which is mutated in 36% of the sequenced clones. *repA* is located on chrII and encodes the replication initiation protein A, which regulates plasmid replication and copy number^41,42^. As mentioned above, chrII is thought to have originated from a plasmid^34^ and relies on the ancestral plasmid genes to replicate and segregate as a chromosome^34,43^. Crucially, the 20 clones that had a mutation in *repA* had increased copy number of the entire chrII (↑chrII), and only five clones with increased copy number of chrII did not have a mutation in *repA* (Fig. 2). Mutations in *repA* are dispersed throughout the entire length of the protein and do not affect any specific domain (Supplementary Fig. 6, Data file 1). Mutations include nine non-synonymous point mutations, a 24 nt insertion, and three deletions that cause frameshifts affecting the C-terminal region. MFA confirmed that despite the mutations in *repA*, chrII replication is unidirectional, as in the wild-type strain (Fig. 1C).

**Figure 2.**
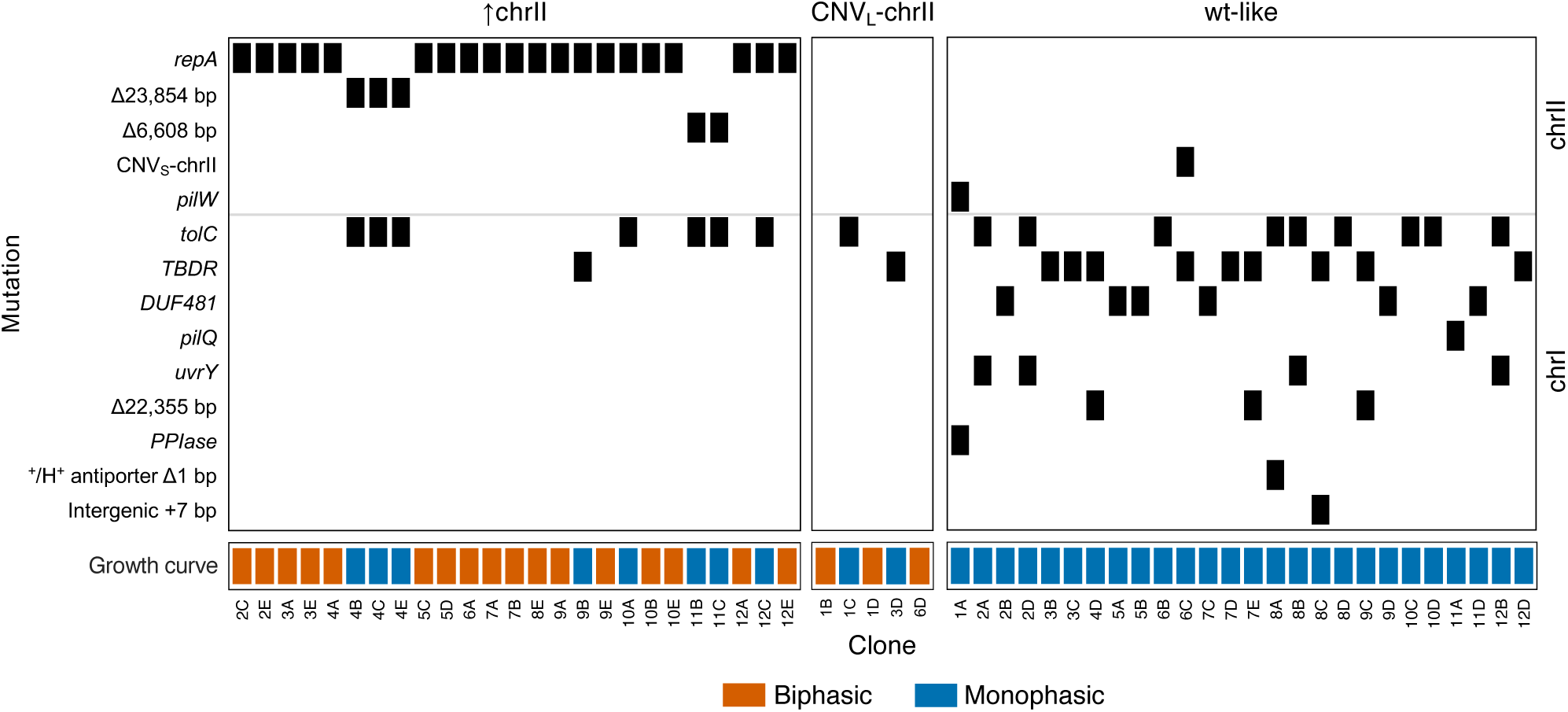
Summary of mutations identified in sequenced clones and the shape of their growth curves. Clones are classified depending on the presence/absence of increases in the copy number of the entire chrII (↑chrII) and CNV_L_-chrII. Orange denotes clones with a biphasic growth curve and blue indicates clones with monophasic growth curves. TBDR, TonB-dependent receptor; DUF481, domain of unknown function 481; PPIase, peptidylprolyl isomerase.

We closely inspected the five clones that had increased copy number of the entire chrII (↑chrII) but no mutations in *repA*. All five clones have one of two types of deletion in chrII, in a region that partially overlaps with a genomic island. Three clones isolated from population 4 have a deletion of 23,854 bp (starting at position 25,864 bp) and two clones from population 11 have a deletion of 6,608 bp (starting at position 46,491 bp). Both deletions overlap in a region spanning four genes (46,491 until 49,718), one of which encodes a putative helicase (PSHA_RS15050) (Supplementary Fig. 7). Helicases unwind double-stranded DNA during replication^44^, and we hypothesize that the truncation or absence of PSHA_RS15050 causes an imbalance in the number of chrII copies, by a mechanism yet to be determined.

In conclusion, we found that all clones that had an increased copy number of the entire chrII (↑chrII) have mutations in either the *repA* gene or a deletion affecting a putative helicase, suggesting that these are responsible for the increase in the number of copies of chrII.

### Mutations in outer membrane proteins are involved in nutrient stress adaptation

Approximately half of the sequenced clones (46%) do not have copy number changes affecting chrII, indicating the existence of alternative pathways for adapting to nutrient stress. To identify these pathways, we closely looked at genes mutated in parallel across multiple populations (Fig. 2). The second most mutated gene in our experiment is *tolC*, mutated in 30% of the sequenced clones. TolC is an outer membrane protein that forms part of a tripartite efflux system involved in the export of antibiotics and other toxic compounds out of the cell^45,46^. Two other outer membrane proteins, TonB-dependent receptor (TBDR) and DUF481, are the third and the fourth most mutated genes in our experiment respectively. TBDR is mutated in 20% of the sequenced clones. It is involved in the uptake of large substrates that do not easily diffuse into the cytoplasm, such as iron-siderophore complexes and vitamin B12^47^. DUF481, mutated in 11% of the clones, belongs to a Pfam clan (CL0193) that groups families of beta-barrel membrane proteins, including porins and TonB-dependent receptors^48^. This suggests that DUF481 might also be an outer membrane protein.

All but two clones have copy number changes in chrII (↑chrII or CNV_L_-chrII) or mutations in tolC, TBDR or DUF481. One of these two clones has a mutation in the *pilQ* gene, and the other has a mutation in the *pilW* gene. PilQ forms a large pore in the outer membrane upon oligomerization and it is involved in DNA uptake, and in the entry of heme and antimicrobial agents^49–51^. PilW stabilizes PilQ and it is required for the formation of PilQ channels^52^.

Disruption of *tolC* in *E. coli* impairs growth on efflux substrates such as fusidic acid and novobiocin^53^. Similarly, mutations affecting *TBDR* result in a lack of active substrate uptake, which improves bacterial survival in toxic compounds^54–56^. To investigate the effect of mutations in *tolC* and *TBDR*, we grew *P. haloplanktis* TAC125 wild-type clone and clones carrying only mutations in *tolC* or *TBDR* in fusidic acid. As all clones carrying mutations in *tolC*, *TBDR*, *DUF481*, *pilQ* and *pilW* exhibit a monophasic growth curve (Fig. 2, Supplementary Fig. 8), we also assessed the growth of clones with point mutations in *DUF481*, *pilQ*, and *pilW* in the presence of fusidic acid. While the wild-type clone grows well in the presence of fusidic acid, clones with mutations in these five genes barely grow in this antibiotic (Fig. 3). This suggests that mutations in these genes impair the ability to export toxic compounds.

**Figure 3.**
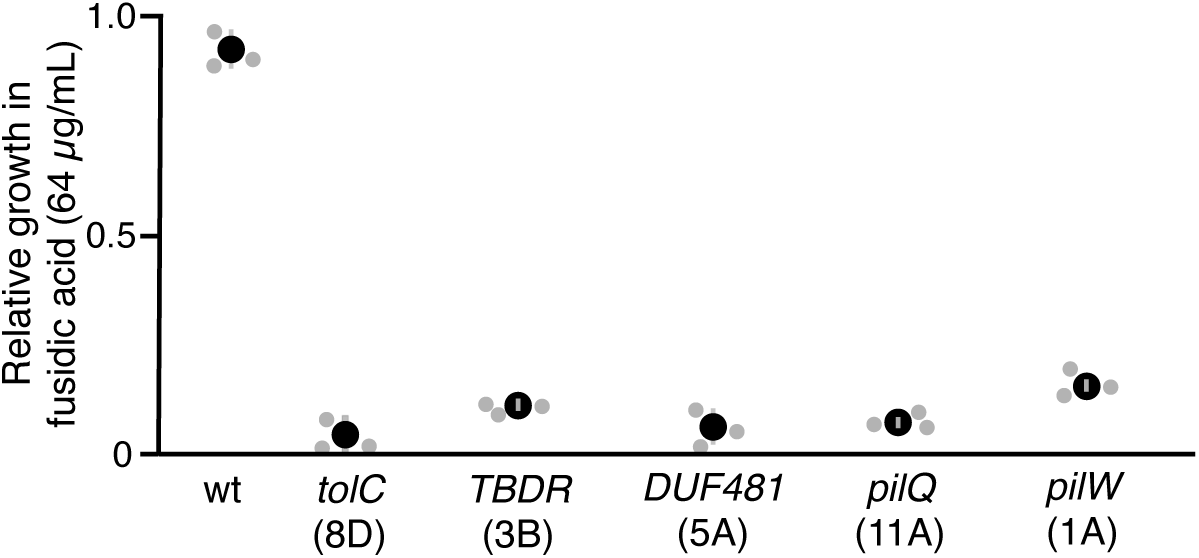
Growth in fusidic acid is impaired by mutations in *tolC*, *TBDR*, *DUF481*, *pilW* and *pilQ*. Mutations in three outer membrane protein genes and two type IV pili genes impact growth in fusidic acid. The plot shows the results for three independent biological replicates normalized by a treatment without fusidic acid. Black circles indicate the mean relative growth. Bars represent the standard error (SE). wt, wild-type; TBDR, TonB-dependent receptor; DUF481, domain of unknown function 481. Clone 1A has an additional mutation in a gene coding for a peptidylprolyl isomerase (PPIase).

Our results show that mutations in five genes (*tolC*, *TBDR*, *DUF481*, *pilQ* and *pilW*), potentially encoding for outer membrane proteins involved in the transport of various compounds, facilitate adaptation to gluconate limitation. Mutations in these genes are mutually exclusive, suggesting that these are convergent ways of coping with the severe nutrient stress. Further experiments are required to elucidate the molecular mechanism underlying the mutations in the outer membrane proteins.

### CNVs in chrII are an unstable mechanism for adapting to nutrient stress

The most frequent copy number change in our experiment is the increase in the copy number of the entire chrII (↑chrII). However, as previously mentioned, there are five clones, isolated from three different endpoint populations, that have a CNV affecting a large genomic region in chrII (CNV_L_-chrII) (Fig. 2). Three of these clones do not have any additional mutations (Fig. 2). CNV_L_-chrII always comprises the region from position 132,527 to 619,758, which represents 77% of the chrII and includes the operon involved in the transport and metabolism of D-gluconic acid (Fig. 1B and 4A). A CNV that covers almost 77% of the chrII is remarkably long. We hypothesized that it could be a self-replicating circular extrachromosomal DNA, as previously reported in yeast growing in nutrient-limited conditions^57^. However, the amplified region lacks the origin of replication of chrII and most of the genes involved in plasmid partitioning and segregation (*repA*, *parA*, *parB* and *tus*) (Fig. 4A). In addition, the MFA indicated that there are no unannotated additional origins of replication on chrII and that the amplified region is not integrated into chrI (Fig. 1C). These results suggest that CNV_L_-chrII is the consequence of a tandem duplication, a conclusion that is also supported by long-read Oxford Nanopore sequencing (Supplementary Fig. 9).

**Figure 4.**
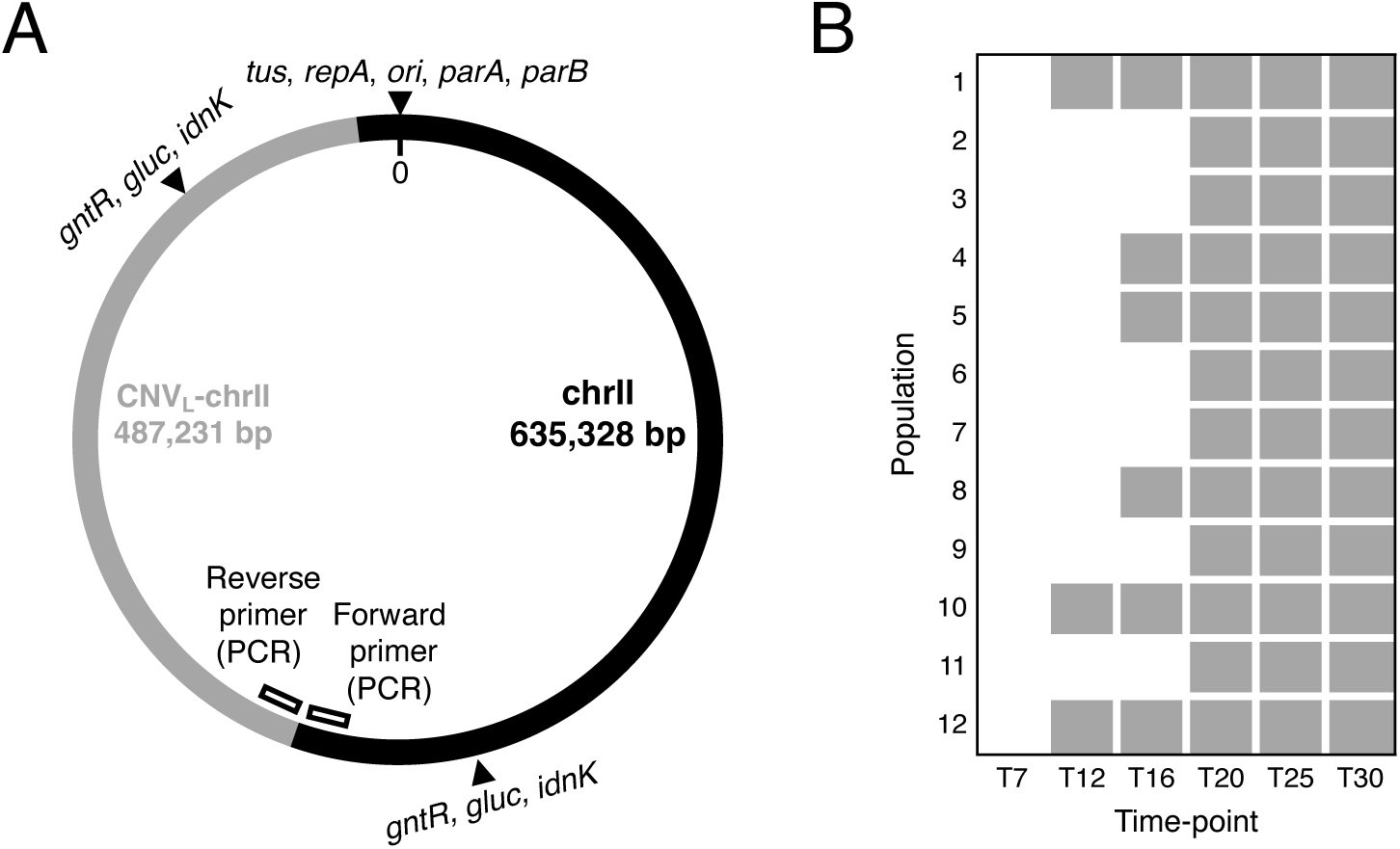
CNV_L_-chrII and its emergence during adaptation to nutrient stress. A: Schematic representation of a chrII copy harboring a CNV_L_-chrII as a large tandem duplication. The standard chrII copy is depicted in black and the CNV_L_-chrII in gray. Black rectangles symbolize the set of primers used for PCR to detect the new junction region generated by the insertion of the CNV_L_-chrII. Black triangles indicate the position of genes involved in the replication, partitioning and segregation of chrII (*tus*, *repA*, *ori*, *parA*, *parB*), and also the locations of the gluconate operon (*gntR*, *gluc*, *idnK*). B: Emergence of CNV_L_-chrII across revived populations and time-points in the evolution experiment. The presence of CNV_L_-chrII was assessed by PCR via the amplification of the newly generated junction between the standard chrII and the CNV_L_-chrII.

To investigate the emergence and dynamics of CNV_L_-chrII, we examined its presence in populations revived from intermediate time points of our experiment. We designed PCR primers to amplify the new junction region at the intersection between the first and the second copy of the tandem duplication (Fig. 4A). We first detected CNV_L_-chrII at the population level in transfer 12, and its frequency increased thereafter. By transfer 20, we detected it in all populations (Fig. 4B). Despite this, only five out of the 56 sequenced endpoint clones had the CNV_L_-chrII, indicating that it may be unstable (Fig. 1A and 1B). A further line of evidence suggests this instability. After isolating colonies obtained from streaking a clone carrying a CNV_L_-chrII, not all the single colonies had the CNV_L_-chrII. The rate of CNV_L_-chrII loss fluctuated around 25% over three replicates (Supplementary Table 3).

In addition to CNV_L_-chrII, we identified a shorter CNV (CNV_S_-chrII) in a single clone (6C). This CNV spans 11% of chrII (from position 539,869 to 609,278) (Fig. 2, Data file 1). It starts in the middle of the gluconate transcriptional repressor (PSHA_RS17140) and includes the gluconate transporter (PSHA_RS17145) and the gluconokinase (PSHA_RS17150) genes, providing further evidence for the beneficial effect of the increased expression of the gluconate operon in our experimental setup.

Taken together, our results show that CNV_L_-chrII is rapidly gained in response to stress, but it is unstable. Both CNV_L_-chrII and CNV_S_-chrII have a region of microhomology (six or seven nucleotides) at the beginning and end of the CNV (Supplementary Fig. 10). These sites appear to recombine specifically, reversibly, and at high frequency.

### Increases in the copy number of chrII provide a fitness advantage

Our study identified two evolutionary pathways to adapt to nutrient stress. One involves copy number changes that affect chrII (↑chrII or CNV_L_-chrII, 54% of clones), and the other involves mutations in genes encoding outer membrane proteins (64% of clones). They act on different molecular pathways and have very different phenotypic effects. After 200 generations of evolution, neither of the two evolutionary pathways became fixed in any of the 12 endpoint populations. Instead, they coexisted in endpoint populations, and in some cases, even in clones (Fig. 2).

We assessed the fitness of the two evolutionary pathways using growth curves. Clones with chrII copy number changes (either ↑chrII or CNV_L_-chrII) and no other mutations have a biphasic growth curve similar to the wild-type strain, but the lag phase is shorter, and the first stationary phase reaches a higher bacterial density (Fig. 5). As previously stated, this increase in growth is likely mediated by a gene dosage effect due to a higher expression of the gluconate operon located in chrII.

**Figure 5.**
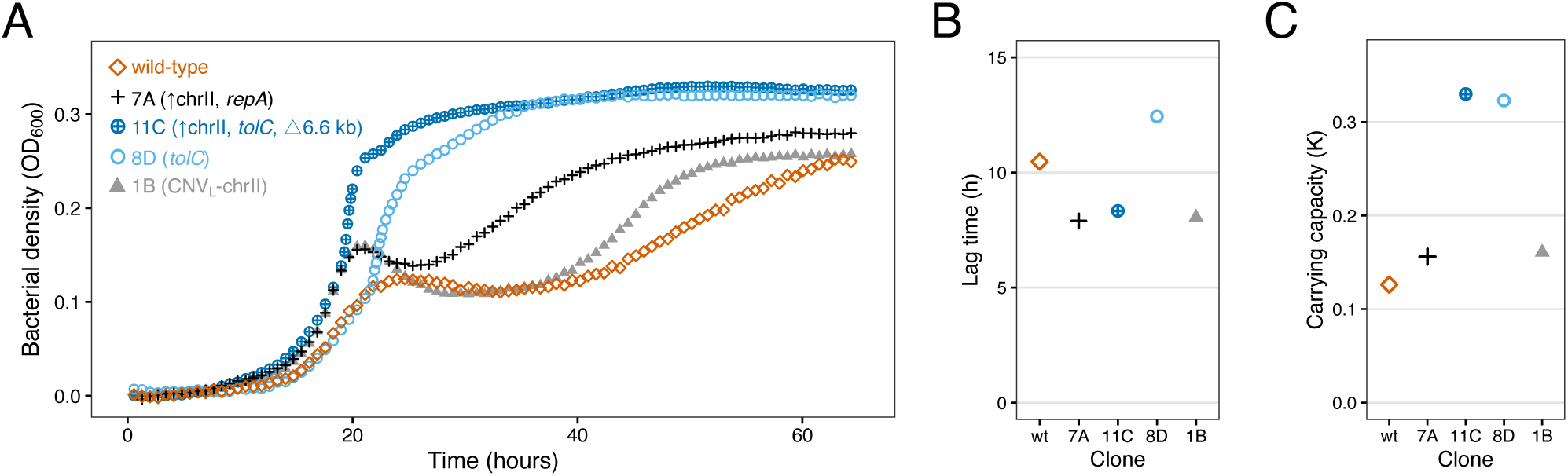
Impact on bacterial fitness of copy number changes affecting chrII and mutations in genes encoding outer membrane proteins. A: Growth comparison between the wild-type clone and clones adapted to nutrient stress. Growth curves represent the average of three technical replicates. TBDR, TonB-dependent receptor; DUF481, domain of unknown function 481. B: Comparison of lag phase duration among selected clones. Lag phase duration was determined as the time when the biomass overcomes the minimal detectable threshold, which corresponds to OD_600_=0.01 (absorbance at 600 nm). C: Comparison of the carrying capacity among selected clones. Carrying capacity was quantified during the first growth phase.

Clones carrying mutations only in a gene coding for an outer membrane protein (*tolC*, *TBDR*, *DUF481, pilQ*), also show an increase in fitness relative to the wild-type strain. These clones have a monophasic growth curve and reach higher bacterial density levels than the wild type (Fig. 5, Supplementary Fig. 8). Mutations in these outer membrane proteins hinder the export of fusidic acid (Fig. 3), suggesting that impaired transport of an unknown compound may facilitate growth in our nutrient-poor medium.

Clones following either evolutionary pathway showed improved growth compared to the wild-type strain. These two pathways are not mutually exclusive, and a few clones (10) carry both (Fig. 2, Fig. 5). Clones carrying both an increased copy number of chrII and mutations in one of the outer membrane proteins are the fittest. These clones show the growth advantages associated with both pathways: a shortened lag phase and increased bacterial density (Fig. 5).

## DISCUSSION

Aneuploidies and CNVs are well known drivers of rapid adaptation to stress in eukaryotes^1,4–11,29^. In bacteria, adaptive CNVs are also prevalent, but the amplified region is usually limited to a gene or a genomic region^15,17–19,58^ and the gain of an entire chromosome has not been reported. In this study, we investigated adaptation to nutrient stress in a bacterium with two chromosomes (chrI and chrII) and uncovered a new mechanism for stress adaptation that is analogous to aneuploidy in eukaryotes. As approximately 10% of bacterial species, including some important pathogens, have a multipartite genome^31,32,59^, this new mechanism may be a relevant adaptive mechanism that has been overlooked.

In our experiment, increases in the copy number of the entire chrII (↑chrII) dominated the rapid adaptation of *P. haloplanktis* TAC125 to nutrient stress. Most clones with increased copy number of the entire chrII had mutations in the *repA* gene, a remnant of the ancestral plasmid that gave rise to chrII^34,37^. In plasmids, RepA initiates DNA replication and regulates plasmid copy number^60,61^ and mutations in *repA* have been associated with changes in plasmid copy number^42,62^. We therefore hypothesize that the *repA* mutations detected in our experiment are driving the increase in the copy number of chrII.

Compared to yeast and *C. albicans*, where CNVs appear to be more common than aneuploidy during short-term stress adaptation^9,10^, we only identified six CNVs in chrII among the sequenced clones. Five CNVs (CNV_L_-chrII) start and end at the same exact genomic coordinates. The remaining CNV is shorter (CNV_S_-chrII) and partially overlaps with CNV_L_-chrII. The lack of diversity and high degree of repeatability suggests a common mechanism of origin for the observed CNVs. In contrast, CNVs in yeast and *C. albicans* are diverse and rarely occur in the exact same genomic location, indicating that they are likely formed through different mechanisms^9^. Both CNV_L_-chrII and CNV_S_-chrII are tandem direct duplications, but there are no insertion sequences (IS), inverted repeats or repetitive extragenic palindromic (REP) sequences in the proximity that could explain their origin. However, a genomic island containing a tyrosine-type recombinase/integrase is predicted to overlap with the end of both CNV_L_-chrII and CNV_S_-chrII (from position 609456 to 628865). Additionally, at the extremes of the duplicated region in both CNVs there is a region of DNA microhomology consisting of six or seven nucleotides, and these nucleotides are found at the junction site (Supplementary Fig. 10). Short and long read sequencing revealed that no additional nucleotides were inserted at the novel junction site. One possibility is that both CNVs have originated through site-specific illegitimate recombination mediated by an end-joining mechanism to repair a DNA break^63–66^. However, it is noteworthy that in CNV_L_-chrII the short homology regions are located almost 0.5 Mb apart.

Most clones with copy number changes in chrII (↑chrII, CNV_L_-chrII) have a copy number between 1.25 and two (Fig. 1A, Supplementary Fig. S4). This is expected for clones with increased copy number of the entire chrII, as different mutations in *repA* will translate into different efficiency in replication initiation. In the case of CNV_L_-chrII, a putative tandem duplication, the likely explanation is that, due to its extraordinary length, the CNV_L_-chrII is unstable and a significant fraction of the cells lose it, resulting in a population with a mixture of clones. In addition, in evolving populations, clones with CNV_L_-chrII are likely to be outcompeted by clones with increased copy number of the entire chrII or mutations in outer membrane proteins, which grow better and have a larger fitness advantage (Fig. 5). However, neither copy number changes affecting chrII nor mutations in outer membrane proteins, or their combination, are fixed in any of the twelve populations. The slow fixation may be caused by clonal interference between clones with different beneficial mutations of varying fitness effects^67^. However, it could also point to a limitation of our experiment, its short duration.

Both types of chrII copy number changes (↑chrII and CNVs), confer a growth advantage probably due to the increased expression of the gluconate operon, which metabolizes D-gluconic acid, the only carbon source present at very low concentrations in our media. Contrary to previous studies^9,18^, changes in the expression of the gluconate operon are driven solely by copy number changes in chrII. We did not find a combination of CNV and point mutations affecting the gluconate operon, nor point mutations in the gluconate operon. It may be that copy number changes dominate the rapid adaptation to D-gluconic scarcity because they are a very frequent type of beneficial mutation, since any copy number change, either mediated by *repA* mutations or CNVs, that contains the gluconate operon is selected for at low D-gluconic concentrations^2,18^.

Copy number changes are costly and unstable and they usually revert if the stress disappears^15^ or are replaced by less costly mutations if the stress persist^2,29^. In contrast to other studies, the two types of copy number change we found differ in their mechanism of origin, which affects their potential for reversibility. CNV_L_-chrII and CNV_S_-chrII are tandem duplications and are completely reversible, leaving no scar in the genome. Conversely, this may not be the case for the increased copy number of the entire chrII. Increases in chrII copy number are mediated by point mutations in *repA* and deletions affecting a putative helicase. Given that these mutations cannot be reversed, additional compensatory mutations may be necessary to restore the chrII copy number to one, which would leave two mutations in the genome. In our experiment, all clones that had a mutation in *repA* had an increased copy number of the entire chrII, suggesting that we do not have cases of transient increases in chrII copy number that have been replaced by less costly beneficial mutations. However, it could be that the copy number changes in chrII were later replaced by point mutations in the gluconate operon, but that we did not observe the whole process because 200 generations were not enough.

In bacteria, increases in the copy number of a secondary chromosomes may be more stable than eukaryotic aneuploidies because secondary chromosomes contain fewer genes involved in core functions than the main chromosome^68^, potentially reducing the cost of having additional copies. In our experiment, clones with increases in the copy number of the entire chrII only grew slightly slower than the wild type in rich media (Supplementary Fig. 11), suggesting that they may be stable if the stress disappears, at least under laboratory conditions. In addition, bacterial secondary chromosomes are enriched in horizontally transferred genes and in genes involved in adaptation to the environment^33,69^. Therefore, gaining additional complete copies of the secondary chromosome may be beneficial in environments other than the stressful one, which might promote their stability. If the increases in the copy number of the entire chrII are stable, they could affect evolutionary dynamics not only in the short term by increasing gene dosage, but also in the long term by increasing the target size for potentially beneficial mutations, thus accelerating adaptation^9,70^. Elucidating the stability of the increases in the copy number of the entire chrII over time is the subject of future work.

To our knowledge, this is the first time that an adaptive increase in the copy number of an entire chromosome has been detected in bacteria. The scarcity of evolution experiments in bacteria with multipartite genomes and their potential reversibility leaving minimal traces in the genome, may have made their detection difficult. In addition to their transient nature, copy number changes can be challenging to detect when sequencing environmental or clinical samples because they can be rapidly lost when cultured in the laboratory on rich media^26,71^. Because secondary chromosomes usually derive from a plasmid and have plasmid-like replication systems^32,33^, it is tempting to speculate that copy number changes mediated by mutations in the replication protein may be a common mechanism of adaptation to stress in bacteria with multipartite genomes. A similar mechanism has been found to increase plasmid copy number in plasmids hosting different replication systems^42,62,70,72,73^ and the artificial overexpression of the replication initiator protein (RctB) increases the number of copies of *Vibrio cholera*’ secondary chromosome^74–76^. However, mutations in the replication protein can also decrease copy number. In a recent study we found that the increase in the number of chrII copies was costly when *P. haloplanktis* was grown at high temperatures, and the copy number was reduced back to one by compensatory mutations in *repA*^38^. In that case, the initial increase in the copy number of chrII was mediated by deletions in a putative helicase and not by mutations in *repA*. Altogether, we have identified 13 mutations in *repA* that increase the copy number of the entire chrII and 19 mutations that decrease it. This high number of mutations in the *repA* gene suggests that chrII copy number can be easily altered in response to stress.

Overall, our study reveals the existence of a novel mechanism for rapid adaptation to stress in a bacterium with a multipartite genome. This mechanism is mediated by an increase in the copy number of the entire secondary chromosome through mutations in the replication protein. We hypothesize that the plasmidic origin of the secondary chromosomes provides a mechanism for rapidly changing their copy number, enhancing genome plasticity and the adaptive potential of multipartite genomes. Given that secondary chromosomes are enriched in genes involved in interacting with the environment, such as antibiotic and metal resistance genes^33,68,69^, their plasticity could be a mechanism for rapid adaptation to antibiotics. Crucially, pathogenic bacteria like *Burkoldheria cenocepacia* and *Vibrio cholera* possess multipartite genomes. Copy number changes, such as aneuploidy and increased copy number of plasmids carrying antibiotic resistance genes, are well-known drivers of drug resistance^70,72,73,77^. Similarly, in bacteria with multipartite genomes, the increase in the copy number of the secondary chromosome could be a simple and rapid mechanism for antibiotic resistance, analogous to aneuploidy in eukaryotes, that has been overlooked until now.

## MATERIALS AND METHODS

### Strains and growth conditions

We purchased *Pseudoalteromonas haloplanktis* TAC125 from the Centre de Ressources Biologiques de l’Institut Pasteur (CIP108707). We grew bacterial cultures in minimal marine sea water medium -MMSW-(10 g/L NaCl, 1 g/L KH_2_PO_4_, 1 g/L NH_4_NO_3_, 0.2 g/L MgSO_4_ • 7H_2_O, 10 mg/L FeSO_4_ • 7H_2_O, 10 mg/L CaCl_2_ • 2H_2_O, pH=7.0)^78^ supplemented with 1 g/L D-gluconic acid at 15 °C and constant shaking at 230 rpm.

### Evolution experiment

To adapt *P. haloplanktis* TAC125 to a minimal medium with low concentrations of D-gluconic acid we conducted a short evolution experiment^38^. Briefly, we evolved 12 replicate populations starting from a single colony isolated from the wild-type strain for 200 generations, at 15 °C and in MMSW supplemented with D-gluconic acid (0.1%) as the only carbon source. 0.1% D-gluconic acid is the minimum concentration that supports growth; a concentration 10 times higher is normally used^78^. Cultures were grown in 48-well plates (P-5ML-48-C, Axygen) containing 2 mL of medium and constant shaking at 400 rpm. Initially, the wild-type strain needed 48 hours to reach the stationary phase and during the first 30 days of the experiment, we diluted the populations 100-fold every 48 hours. After 30 days, cultures grew faster and we changed the dilution regime to daily 100-fold dilutions for a total of 15 additional days, amounting to a total of 200 generations (Supplementary Fig. 1). We preserved samples periodically in 20% glycerol and stored them at −80 °C.

### Whole-genome Illumina sequencing

We selected 10 individual colonies from each endpoint population from agar plates containing MMSW with 0.1% D-gluconic acid. We assessed their fitness by measuring growth curves in 200 μL of MMSW at 15 °C in an Infinite M Nano (Tecan) microplate reader with constant shaking (527 s, amp-freq: 200-250 rpm orbital). We measured absorbance at 600 nm over four days. We observed two types of growth curves: monophasic and biphasic. Monophasic growth curves follow a standard bacterial growth curve with a lag phase, an exponential phase, and a stationary phase; whereas biphasic growth curves have two sequential exponential phases (Supplementary Fig. 2). Based on the frequency of both types of growth curves, we proportionally selected four to five clones from each population for whole genome sequencing. In total we selected 56 clones.

We carried out genomic DNA extraction with the DNeasy Blood & Tissue kit (QIAGEN). We determined gDNA concentration using a Qubit fluorometer dsDNA Broad Range assay (ThermoFisher Scientific) and Nanodrop 8000 (ThermoFisher Scientific) and we assessed the integrity and quality of the gDNA by agarose gel electrophoresis. Library preparation and genome sequencing (Illumina NovaSeq 6000, 150-base pair paired-end reads) were conducted at the Oxford Genomics Centre (United Kingdom).

To identify mutations relative to the wild-type *P. haloplanktis* TAC125 reference genome (chromosome I: NC_007481.1, chromosome II: NC_007482.1, pMEGA plasmid: MN400773, pMtBL plasmid: AJ224742), we analyzed the reads with the computational pipeline *breseq* (v.0.35.5)^79^. *breseq* uses reference-based alignment approaches to predict single-nucleotide mutations, point insertions and deletions, large deletions, and new junctions. We run *breseq* with the alternative mode (-p option) to predict polymorphic mutations. This option allows the detection of low-frequency mutations and it was useful to verify that mutations in clones carrying increases in the copy number of the entire chrII were present in all copies of chrII. We obtained a mean coverage of 684 reads per genomic site for our samples.

*breseq* detected two CNVs in chrII, CNV_S_-chrII and CNV_L_-chrII. To identify the CNV breakpoints we visually inspected the CNVs using the Integrative Genomics Viewer (IGV) (v.2.13.1)^80^, focusing on split reads. Crucially, split reads mapped to each of the ends of the CNVs. In addition, to study the new junction sequence created as a result of the CNV, we assembled the genomes *de novo* using SPAdes (v.3.15.4)^81^ and the Illumina reads. We aligned the contigs we obtained against the reference genome using BLAST (v2.15.0)^82^.

### Estimation of chromosome copy number

To assess the frequency of clones carrying an increase in the number of copies of chrII we used two methods. First, we used samtools idxstats (v.1.2)^83^ to calculate the read coverage for each chromosome and compute the mean read coverage per base pair. To estimate the number of copies of chrII (ratio) we divided the mean read coverage for chrII by the mean read coverage for chrI. We used the ratio of the wild-type clone to normalize the ratios of all remaining clones. We inferred variations in the copy number of chrII when the ratio was over 1. Since the absolute number of reads obtained from chrI and chrII varied across samples, we corrected the ratios by a factor of 0.72 for visualization purposes. 0.72 is the mean difference of reads mapping to chrII relative to the reads mapping to chrI in clones without variations in the copy number of chrII. We used samtools depth -aa (v.1.2)^83^ to obtain the read coverage at each chrII position. We divided the read depth by the 0.72 correction factor and visualized increments in the number of chrII copies by plotting average read depth as a function of chromosome position in windows of 5000 bp.

Second, we used quantitative PCR (qPCR) to estimate the copy number of genes located on chrII and validate the two copy number changes we detected: increases in the copy number of the entire chrII and CNV_L_-chrII. We used a set of four genes, two housekeeping genes located on chrI (*rpoB* and *gyrA*) and two genes on chrII (*parA* and *gluc*) (Supplementary Table 4). Thus, using the *P. haloplanktis* TAC125 wild-type strain as a control, we expect clones to have a higher relative amount of DNA for those genes located in regions/chromosomes that have increased copy number. Primer pairs were designed with Primer3Plus and they were checked by standard PCR using Thermopol Taq polymerase to confirm the amplification of a single band of the desired size^84^. qPCR reactions contained 10 ng/μL gDNA and were performed in triplicates in 96-well plates (Trefflab) or 384-well plates (Trefflab) using the HOT FIREPol EvaGreen qPCR Mix Plus (Solis Biodyne) in a LightCycler 480 instrument (Roche). The qPCR cycling protocol consisted of an initial activation step at 95 °C (12 min), followed by 40 cycles including denaturation at 95 °C (15 s), annealing at 60 °C (20 s), and extension at 72 °C (20 s). A melt curve from 60 °C to 95 °C was recorded after each run with 0.5 °C increments every 15s. We computed Ct values (cycle threshold) using the absolute quantification method with the LightCycler software (v.4.1) by the second derivative maximum method. We used the ΔΔCt method to obtain the relative abundance of genes located in chrII relative to the genes in chrI. Wild-type values served as normalization factor for this purpose.

### Oxford Nanopore Sequencing (ONT)

To investigate whether the large CNV affecting 77% of chrII (CNV_L_-chrII) is a tandem duplication or a self-replicating circular extrachromosomal DNA we performed long-read sequencing followed by *de novo* assembly. Using the DNeasy Blood & Tissue kit (QIAGEN) we extracted genomic DNA from five clones: the wild-type, one clone with a complete increase in the number of copies of chrII (7A), and three clones with CNV_L_-chrII (clones 1B, 1D, 6D). We used a Qubit fluorometer dsDNA Broad Range assay (ThermoFisher Scientific) and Nanodrop 8000 (ThermoFisher Scientific) to assess gDNA concentration and evaluated its integrity by agarose gel electrophoresis. Oxford Nanopore Sequencing was conducted by Plasmidsaurus using PromethION R10.1.4 Flow Cell and Kit 14 chemistry. We obtained a total number of reads ranging from 72076 to 207155 and a coverage that fluctuated between 90X to 266X (Supplementary Table 5). We filtered reads by quality using Filtlong (v0.2.1) with the default parameters^85^. We run Flye (v2.9.2) using parameters specific for high quality ONT reads to obtain a final assembly in contigs^86^. We visualized the assemblies using Bandage (v0.8.1)^87^ and assigned the contigs to the corresponding replicon of *P. haloplaktis* TAC125 using BLAST (v2.15.0) ^82^.

To detect structural variations we aligned ONT Fastq files to the *P. haloplanktis* TAC125 reference genome using NGMLR (-x ont v.0.2.8)^88^. We used samtools (v.1.2) to sort (samtools sort) and index (samtools index) the mapped reads^83^. We used Sniffles (v1.0.6) to identify structural variants^89^ and confirmed the presence of CNV_L_-chrII in samples from clones 1B, 1D and 6D. We used IGV (v.2.13.1)^80^ to visually inspect the CNV_L_-chrII start and end breakpoints. We identified split reads, and all of them mapped to each of the extremes of the CNV. We also selected reads mapping to the CNV_L_-chrII new junction and used BLAST (v2.15.0) to confirm the lack of mutations or sequence misalignments^82^.

### Marker Frequency Analysis (MFA)

We performed MFA to confirm the copy number changes in chrII and investigate the presence of additional origins of replication in chrII. We cultured four clones in rich medium (marine broth 2216 (Sigma)): a clone with increased copy number of the entire chrII (7A), two clones containing CNV_L_-chrII (1B, 6D), and the wild-type clone. We extracted gDNA at both early-exponential and stationary phase using the DNeasy Blood & Tissue kit (QIAGEN). Library preparation and sequencing was conducted at the Functional Genomics Center Zurich (FGCZ). We used a Truseq PCR Free protocol with NovaSeq 6000 (Illumina), 150–base pair (bp) paired-end reads. We analyzed sequencing reads as previously described^39,40^. Reads were mapped with Bowtie 2^90^ using NC_007481.1, NC_007482.1, MN400773, and NZ_AJ224742 as reference sequences. Mapped reads were binned to 1 kbp windows using a Perl script. 150+ bp sequence repeats were identified with R2R^39^ and windows with repeated sequence excluded. We calculated a correction factor for each window from the stationary samples and applied it to the exponentially grown samples.

### Growth in fusidic acid

We evaluated growth in fusidic acid in clones with single mutations in potential outer membrane proteins (*tolC*, TonB-dependent Receptor -TBDR- and Domain of Unknown Function 481 -DUF481-) and proteins of the type IV pili (PilQ and PilW). We grew the selected clones for 72 h in MMSW supplemented with 0.1% D-gluconic acid with constant shaking at 230 rpm in 96-well plates (TPP; 92096). Then we transferred bacterial cultures to medium (MMSW with 0.1% D-gluconic acid) containing 64 μg/mL of fusidic acid with a starting OD_600_ of 0.02 and we grew them for 72 h at 15 °C and constant shaking at 400 rpm. After 72 h we measured absorbance at 600 nm using an Infinite M Nano (Tecan) microplate reader and we computed the relative growth of each clone by normalizing to the growth of the respective clone in medium (MMSW with 0.1% D-gluconic acid) without fusidic acid.

### Assessing CNVs stability

To investigate the presence of CNV_L_-chrII in intermediate populations from the evolution experiment we designed primers to amplify the new junction region between the end of the first copy and the beginning of the second copy (primers iC2_4 and iC2_6) (Supplementary Table 4). We screened the 12 replicate populations of the experiment at five intermediate time-points (T7, T12, T16, T20, T25) and the endpoint (T30). We assessed each population by mixing 40 μL of culture with 70 μL sterile distilled water. We centrifuged samples for 3 min at 4000 rpm, discarded the supernatant and resuspended the cell pellet in 30 μL sterile distilled water. We incubated the samples for 10 min at 95 °C and centrifuged them at 4000 rpm for 5 min. We used 2 μL of the supernatant to prepare the PCR reactions according to the specifications provided by the manufacturer for the GoTaq G2 DNA Polymerase (Promega). We performed PCR in 96-well plates (Starstedt) using a GeneAmp Thermocycler (Applied Biosystems). We used a PCR cycling protocol with an initial activation step at 95 °C (7 min), followed by 35 cycles including denaturation at 95 °C (30 s), annealing at 55 °C (30 s), and extension at 72 °C (20 s). We assessed the presence of PCR products by 1% agarose gel electrophoresis.

To study the stability of CNV_L_-chrII we used standard PCR with primers iC2_4 and iC2_6 to evaluate its loss in minimal medium in clone 1B. We revived and streaked clone 1B on agar plates containing MMSW supplemented with 0.1% D-gluconic acid and incubated the plates at 15 °C for four days. We isolated 31 individual colonies and grew them in 200 μL MMSW with 0.1% D-gluconic acid for 2 days at 15 °C with constant shaking at 400 rpm. We assessed the presence of the CNV_L_-chrII using PCR amplification, as described above, for each of the 31 cultures. We repeated the whole process three times to have three biological replicates (Supplementary Table 3).

### Fitness estimates of evolved clones

We used growth curves to estimate the fitness of the two evolutionary pathways. We selected three clones carrying single mutations affecting the different pathways and one clone that has both. We inoculated the wild-type and evolved clones in 2 mL of MMSW supplemented with 0.1% D-gluconic acid and incubated the cultures at 15 °C for three days with constant shaking (400 rpm). We measured the optical density (OD_600_) and diluted the cultures to an OD_600_ of 0.005. Using an Infinite M Nano (Tecan) microplate reader we measured growth curves at 15 °C with constant shaking (527 s, amp-freq: 200-250 rpm orbital) over 72 h. We assessed fitness and growth kinetics using two parameters: lag phase duration and carrying capacity (K). We obtained growth curve metrics using the gcplyr R package (v1.9.0)^91^. Since some growth curves deviate from the typical monophasic growth curve, lag time was computed based on the biomass increase taking as a reference an OD_600_ of 0.01^92^. 0.01 is the threshold at which the minimal increase in population size is detectable in our cultures. For biphasic growth curves, we computed the carrying capacity as the maximum growth reached during the first growth phase.

## Supporting information

Supplementary_information-Vega-Cabrera_etal

Supplementary_Data1_Vega-Cabrera_etal

## ACKNOWLEDGMENTS

We would like to thank to Álvaro San Millán, Jakub Czarnecki, Marie-Eve Kennedy-Val, Didier Mazel, Ricardo León Sampedro and the Evolutionary Biology and Evolutionary Microbiology groups at ETH Zurich for fruitful discussions. We thank Miriam Olombrada for technical support. We thank Enora Marrec and Melih Çakar for their help with experiments. qPCRs and gDNA quality assessment were performed at the Genetic Diversity Centre (GDC), ETH Zurich. We thank the Oxford Genomics Centre at the Wellcome Centre for Human Genetics (funded by Wellcome Trust grant reference 203141/Z/16/Z) for the generation and initial processing of the sequencing data. We acknowledge the Functional Genomics Center Zurich (UZH/ETHZ) and Plasmidsaurus for the generation of the sequencing data for the MFA analysis and the Oxford Nanopore sequencing data respectively.

This work was funded by the Swiss National Science Foundation, PRIMA grant PR00P3_185899 (to M.T.-R).

## Author contributions

Conceptualization: MTR. Experimental design: LVC and MTR. Experiments performance: LVC. Data analysis: LVC, OS, MTR. Visualization: LVC, OS, MTR. Writing—original draft: LVC and MTR. Writing—review and editing: LVC, OS, MTR Sequence reads are available in the European Nucleotide Archive database under the accession code XXX.

## REFERENCES

1. Berman, J. Ploidy plasticity: a rapid and reversible strategy for adaptation to stress. FEMS Yeast Res. 16, fow020 (2016).

2. Yona, A. H., Frumkin, I. & Pilpel, Y. A relay race on the evolutionary adaptation spectrum. Cell 163, 549–559 (2015).

3. Tang, Y.-C. & Amon, A. Gene copy-number alterations: a cost-benefit analysis. Cell 152, 394–405 (2013).

4. Anderson, R. P. & Roth, J. R. Tandem genetic duplications in phage and bacteria. Annu. Rev. Microbiol. 31, 473–505 (1977).

5. Beroukhim, R. et al. The landscape of somatic copy-number alteration across human cancers. Nature 463, 899–905 (2010).

6. Dulmage, K. A., Darnell, C. L., Vreugdenhil, A. & Schmid, A. K. Copy number variation is associated with gene expression change in archaea. *Microb*. Genomics 4, e000210 (2018).

7. Zarrei, M., MacDonald, J. R., Merico, D. & Scherer, S. W. A copy number variation map of the human genome. Nat. Rev. Genet. 16, 172–183 (2015).

8. Dolatabadian, A., Patel, D. A., Edwards, D. & Batley, J. Copy number variation and disease resistance in plants. Theor. Appl. Genet. 130, 2479–2490 (2017).

9. Lauer, S. et al. Single-cell copy number variant detection reveals the dynamics and diversity of adaptation. PLoS Biol. 16, e3000069 (2018).

10. Todd, R. T. & Selmecki, A. Expandable and reversible copy number amplification drives rapid adaptation to antifungal drugs. Elife 9, e58349 (2020).

11. Kondrashov, F. A. Gene duplication as a mechanism of genomic adaptation to a changing environment. Proc. R. Soc. B Biol. Sci. 279, 5048–5057 (2012).

12. Brown, C. J., Todd, K. M. & Rosenzweig, R. F. Multiple duplications of yeast hexose transport genes in response to selection in a glucose-limited environment. Mol. Biol. Evol. 15, 931–942 (1998).

13. Gresham, D. et al. The repertoire and dynamics of evolutionary adaptations to controlled nutrient-limited environments in yeast. PLoS Genet. 4, e1000303 (2008).

14. Ben-David, U. & Amon, A. Context is everything: aneuploidy in cancer. Nat. Rev. Genet. 21, 44–62 (2020).

15. Andersson, D. I. & Hughes, D. Gene amplification and adaptive evolution in bacteria. Annu. Rev. Genet. 43, 167–195 (2009).

16. Nicoloff, H., Hjort, K., Levin, B. R. & Andersson, D. I. The high prevalence of antibiotic heteroresistance in pathogenic bacteria is mainly caused by gene amplification. Nat. Microbiol. 4, 504–514 (2019).

17. Tomanek, I. et al. Gene amplification as a form of population-level gene expression regulation. *Nat*. Ecol. Evol. 4, 612–625 (2020).

18. Tomanek, I. & Guet, C. C. Adaptation dynamics between copy-number and point mutations. Elife 11, e82240 (2022).

19. Blount, Z. D. et al. Genomic and phenotypic evolution of *Escherichia coli* in a novel citrate-only resource environment. Elife 9, e55414 (2020).

20. Belikova, D., Jochim, A., Power, J., Holden, M. T. & Heilbronner, S. “Gene accordions” cause genotypic and phenotypic heterogeneity in clonal populations of *Staphylococcus aureus*. Nat. Commun. 11, 3526 (2020).

21. Siegel, J. J. & Amon, A. New insights into the troubles of aneuploidy. Annu. Rev. Cell Dev. Biol. 28, 189–214 (2012).

22. Sheltzer, J. M. et al. Aneuploidy drives genomic instability in yeast. Science 333, 1026– 1030 (2011).

23. Sheltzer, J. M. & Amon, A. The aneuploidy paradox: costs and benefits of an incorrect karyotype. Trends Genet. 27, 446–453 (2011).

24. Torres, E. M. et al. Effects of aneuploidy on cellular physiology and cell division in haploid yeast. science 317, 916–924 (2007).

25. Sunshine, A. B. et al. Aneuploidy shortens replicative lifespan in *Saccharomyces cerevisiae*. Aging Cell 15, 317–324 (2016).

26. Gilchrist, C. & Stelkens, R. Aneuploidy in yeast: Segregation error or adaptation mechanism? Yeast 36, 525–539 (2019).

27. Chen, G., Bradford, W. D., Seidel, C. W. & Li, R. Hsp90 stress potentiates rapid cellular adaptation through induction of aneuploidy. Nature 482, 246–250 (2012).

28. Selmecki, A. M., Dulmage, K., Cowen, L. E., Anderson, J. B. & Berman, J. Acquisition of aneuploidy provides increased fitness during the evolution of antifungal drug resistance. PLoS Genet. 5, e1000705 (2009).

29. Yona, A. H. et al. Chromosomal duplication is a transient evolutionary solution to stress. Proc. Natl. Acad. Sci. 109, 21010–21015 (2012).

30. Adler, M., Anjum, M., Berg, O. G., Andersson, D. I. & Sandegren, L. High fitness costs and instability of gene duplications reduce rates of evolution of new genes by duplication-divergence mechanisms. Mol. Biol. Evol. 31, 1526–1535 (2014).

31. Dicenzo, G. C. & Finan, T. M. The divided bacterial genome: structure, function, and evolution. Microbiol. Mol. Biol. Rev. 81, 10–1128 (2017).

32. Harrison, P. W., Lower, R. P., Kim, N. K. & Young, J. P. W. Introducing the bacterial ‘chromid’: not a chromosome, not a plasmid. Trends Microbiol. 18, 141–148 (2010).

33. Dicenzo, G. C., Mengoni, A. & Perrin, E. Chromids aid genome expansion and functional diversification in the family *Burkholderiaceae*. Mol. Biol. Evol. 36, 562–574 (2019).

34. Médigue, C. et al. Coping with cold: the genome of the versatile marine Antarctica bacterium *Pseudoalteromonas haloplanktis* TAC125. Genome Res. 15, 1325–1335 (2005).

35. Qi, W. et al. New insights on *Pseudoalteromonas haloplanktis* TAC125 genome organization and benchmarks of genome assembly applications using next and third generation sequencing technologies. Sci. Rep. 9, 16444 (2019).

36. Liao, L. et al. Multipartite genomes and the sRNome in response to temperature stress of an Arctic *Pseudoalteromonas fuliginea* BSW20308. Environ. Microbiol. 21, 272–285 (2019).

37. Xie, B.-B. et al. Evolutionary trajectory of the replication mode of bacterial replicons. MBio 12, 10–1128 (2021).

38. Toll-Riera, M., Olombrada, M., Castro-Giner, F. & Wagner, A. A limit on the evolutionary rescue of an Antarctic bacterium from rising temperatures. Sci. Adv. 8, eabk3511 (2022).

39. Skovgaard, O., Bak, M., Løbner-Olesen, A. & Tommerup, N. Genome-wide detection of chromosomal rearrangements, indels, and mutations in circular chromosomes by short read sequencing. Genome Res. 21, 1388–1393 (2011).

40. Val, M.-E. et al. A checkpoint control orchestrates the replication of the two chromosomes of *Vibrio cholerae*. Sci. Adv. 2, e1501914 (2016).

41. Xia, G., Manen, D., Yu, Y. & Caro, L. In vivo and in vitro studies of a copy number mutation of the RepA replication protein of plasmid pSC101. J. Bacteriol. 175, 4165–4175 (1993).

42. Thompson, M. G. et al. Isolation and characterization of novel mutations in the pSC101 origin that increase copy number. Sci. Rep. 8, 1590 (2018).

43. Sonnenberg, C. B. & Haugen, P. The Pseudoalteromonas multipartite genome: distribution and expression of pangene categories, and a hypothesis for the origin and evolution of the chromid. G3 11, jkab256 (2021).

44. Konieczny, I. Strategies for helicase recruitment and loading in bacteria. EMBO Rep. 4, 37–41 (2003).

45. Morona, R., Manning, P. & Reeves, P. Identification and characterization of the TolC protein, an outer membrane protein from *Escherichia coli*. J. Bacteriol. 153, 693–699 (1983).

46. Piddock, L. J. Multidrug-resistance efflux pumps? not just for resistance. Nat. Rev. Microbiol. 4, 629–636 (2006).

47. Schauer, K., Rodionov, D. A. & de Reuse, H. New substrates for TonB-dependent transport: do we only see the ‘tip of the iceberg’? Trends Biochem. Sci. 33, 330–338 (2008).

48. Mistry, J. et al. Pfam: The protein families database in 2021. Nucleic Acids Res. 49, D412–D419 (2021).

49. Craig, L., Forest, K. T. & Maier, B. Type IV pili: dynamics, biophysics and functional consequences. Nat. Rev. Microbiol. 17, 429–440 (2019).

50. Nandi, S., Swanson, S., Tomberg, J. & Nicholas, R. A. Diffusion of antibiotics through the PilQ secretin in *Neisseria gonorrhoeae* occurs through the immature, sodium dodecyl sulfate-labile form. J. Bacteriol. 197, 1308–1321 (2015).

51. Weaver, S. J. et al. CryoEM structure of the type IVa pilus secretin required for natural competence in *Vibrio cholerae*. Nat. Commun. 11, 5080 (2020).

52. Yaman, D. & Averhoff, B. Functional dissection of structural regions of the *Thermus thermophilus* competence protein PilW: Implication in secretin complex stability, natural transformation and pilus functions. Biochim. Biophys. Acta BBA-Biomembr. 1863, 183666 (2021).

53. Touzé, T. et al. Interactions underlying assembly of the *Escherichia coli* AcrAB–TolC multidrug efflux system. Mol. Microbiol. 53, 697–706 (2004).

54. Kadner, R. J., Heller, K., Coulton, J. W. & Braun, V. Genetic control of hydroxamate-mediated iron uptake in *Escherichia coli*. J. Bacteriol. 143, 256–264 (1980).

55. Pugsley, A., Zimmerman, W. & Wehrli, W. Highly efficient uptake of a rifamycin derivative via the FhuA-TonB-dependent uptake route in *Escherichia coli*. J. Gen. Microbiol. 133, 3505–3511 (1987).

56. Silale, A. & van den Berg, B. TonB-dependent transport across the bacterial outer membrane. Annu. Rev. Microbiol. 77, 67–88 (2023).

57. Gresham, D. et al. Adaptation to diverse nitrogen-limited environments by deletion or extrachromosomal element formation of the GAP1 locus. Proc. Natl. Acad. Sci. 107, 18551–18556 (2010).

58. Reams, A. B. & Roth, J. R. Mechanisms of gene duplication and amplification. Cold Spring Harb. Perspect. Biol. 7, a016592 (2015).

59. Ramachandran, R., Jha, J. & Chattoraj, D. K. Chromosome segregation in *Vibrio cholerae*. J. Mol. Microbiol. Biotechnol. 24, 360–370 (2015).

60. Nordström, K. Plasmid R1—replication and its control. Plasmid 55, 1–26 (2006).

61. Ehrenberg, M. & Sverredal, A. A model for copy number control of the plasmid R1. J. Mol. Biol. 246, 472–485 (1995).

62. Hashiro, S. & Yasueda, H. Plasmid copy number mutation in repA gene encoding RepA replication initiator of cryptic plasmid pHM1519 in *Corynebacterium glutamicum*. Biosci. Biotechnol. Biochem. 82, 2212–2224 (2018).

63. Darmon, E. & Leach, D. R. Bacterial genome instability. Microbiol. Mol. Biol. Rev. 78, 1– 39 (2014).

64. Reams, A. B. & Neidle, E. L. Gene amplification involves site-specific short homology-independent illegitimate recombination in *Acinetobacter* sp. strain ADP1. J. Mol. Biol. 338, 643–656 (2004).

65. Ottaviani, D., LeCain, M. & Sheer, D. The role of microhomology in genomic structural variation. Trends Genet. 30, 85–94 (2014).

66. Reams, A. B. & Neidle, E. L. Genome plasticity in *Acinetobacter*: new degradative capabilities acquired by the spontaneous amplification of large chromosomal segments. Mol. Microbiol. 47, 1291–1304 (2003).

67. Gerrish, P. J. & Lenski, R. E. The fate of competing beneficial mutations in an asexual population. Genetica 102, 127–144 (1998).

68. Dicenzo, G. C. & Finan, T. M. Genetic redundancy is prevalent within the 6.7 Mb *Sinorhizobium meliloti* genome. Mol. Genet. Genomics 290, 1345–1356 (2015).

69. Riccardi, C. et al. Independent origins and evolution of the secondary replicons of the class Gammaproteobacteria. *Microb*. Genomics 9, 001025 (2023).

70. San Millan, A., Escudero, J. A., Gifford, D. R., Mazel, D. & MacLean, R. C. Multicopy plasmids potentiate the evolution of antibiotic resistance in bacteria. *Nat*. Ecol. Evol. 1, 0010 (2016).

71. Sandegren, L. & Andersson, D. I. Bacterial gene amplification: implications for the evolution of antibiotic resistance. Nat. Rev. Microbiol. 7, 578–588 (2009).

72. DelaFuente, J. et al. Within-patient evolution of plasmid-mediated antimicrobial resistance. *Nat*. Ecol. Evol. 6, 1980–1991 (2022).

73. Dimitriu, T., Matthews, A. C. & Buckling, A. Increased copy number couples the evolution of plasmid horizontal transmission and plasmid-encoded antibiotic resistance. Proc. Natl. Acad. Sci. 118, e2107818118 (2021).

74. Koch, B., Ma, X. & Løbner-Olesen, A. rctB mutations that increase copy number of *Vibrio cholerae* oriCII in *Escherichia coli*. Plasmid 68, 159–169 (2012).

75. Kothapalli, R. et al. The dimerization interface of initiator RctB governs chaperone and enhancer dependence of *Vibrio cholerae* chromosome 2 replication. Nucleic Acids Res. 50, 4529–4544 (2022).

76. Srivastava, P. & Chattoraj, D. K. Selective chromosome amplification in *Vibrio cholerae*. Mol. Microbiol. 66, 1016–1028 (2007).

77. Todd, R. T., Wikoff, T. D., Forche, A. & Selmecki, A. Genome plasticity in *Candida albicans* is driven by long repeat sequences. Elife 8, e45954 (2019).

78. Sannino, F. et al. A novel synthetic medium and expression system for subzero growth and recombinant protein production in *Pseudoalteromonas haloplanktis* TAC125. Appl. Microbiol. Biotechnol. 101, 725–734 (2017).

79. Deatherage, D. E. & Barrick, J. E. Identification of mutations in laboratory-evolved microbes from next-generation sequencing data using breseq. Eng. Anal. Multicell. Syst. Methods Protoc. 165–188 (2014).

80. Robinson, J. T. et al. Integrative genomics viewer. Nat. Biotechnol. 29, 24–26 (2011).

81. Bankevich, A. et al. SPAdes: a new genome assembly algorithm and its applications to single-cell sequencing. J. Comput. Biol. 19, 455–477 (2012).

82. Altschul, S. F., Gish, W., Miller, W., Myers, E. W. & Lipman, D. J. Basic local alignment search tool. J. Mol. Biol. 215, 403–410 (1990).

83. Danecek, P. et al. Twelve years of SAMtools and BCFtools. Gigascience 10, giab008 (2021).

84. Untergasser, A. et al. Primer3—new capabilities and interfaces. Nucleic Acids Res. 40, e115–e115 (2012).

85. 85. Wick, R. R. & Menzel, P. Filtlong. Available Online Github ComrrwickFiltlong Accessed 15 August 2021 (2017).

86. Kolmogorov, M., Yuan, J., Lin, Y. & Pevzner, P. A. Assembly of long, error-prone reads using repeat graphs. Nat. Biotechnol. 37, 540–546 (2019).

87. Wick, R. R., Schultz, M. B., Zobel, J. & Holt, K. E. Bandage: interactive visualization of de novo genome assemblies. Bioinformatics 31, 3350–3352 (2015).

88. Sedlazeck, F. J. et al. Accurate detection of complex structural variations using single-molecule sequencing. Nat. Methods 15, 461–468 (2018).

89. Smolka, M. et al. Detection of mosaic and population-level structural variants with Sniffles2. Nat. Biotechnol. 1–10 (2024).

90. Langmead, B. & Salzberg, S. L. Fast gapped-read alignment with Bowtie 2. Nat. Methods 9, 357–359 (2012).

91. Blazanin, M. gcplyr: an R package for microbial growth curve data analysis. BMC Bioinformatics 25, 232 (2024).

92. Smug, B. J., Opalek, M., Necki, M. & Wloch-Salamon, D. Microbial lag calculator: A shiny-based application and an R package for calculating the duration of microbial lag phase. Methods Ecol. Evol. 15, 301–307 (2024).

93. Varadi, M., et al. AlphaFold Protein Structure Database: massively expanding the structural coverage of protein-sequence space with high-accuracy models. Nucleic acids research 50, D439–D444 (2022).

94. Land, H., & Humble, M. S. YASARA: a tool to obtain structural guidance in biocatalytic investigations. Protein engineering: methods and protocols, 43–67 (2018).

