## Supplementary_information-Vega-Cabrera_etal for "Chromosomal plasticity can drive rapid adaptation in bacteria"

[1]Institute of Integrative Biology, ETH Zurich, Zurich, Switzerland
[2]Department of Science and Environment, Roskilde University, Roskilde, Denmark
‡Present address: Institut de Biologia Evolutiva (CSIC-UPF), Barcelona, Spain

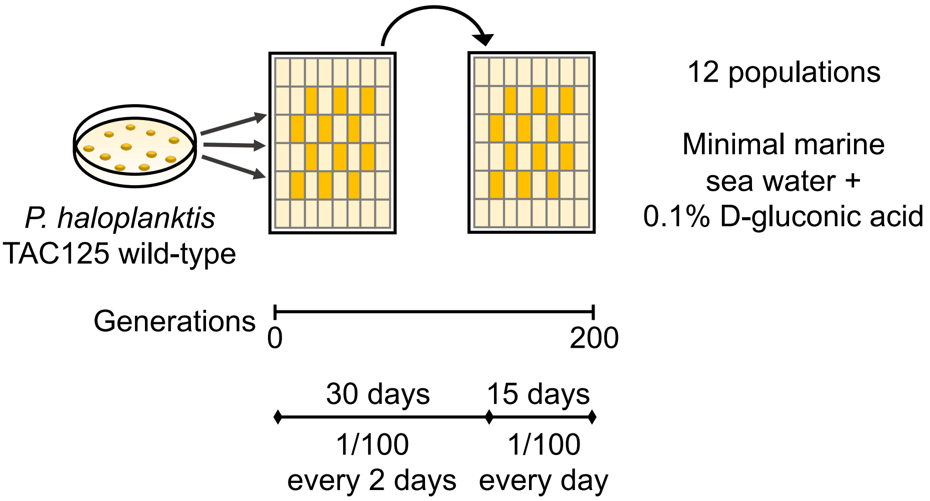


Supplementary Figure 1. Schematic representation of the evolution experiment to adapt *Pseudoalteromonas haloplanktis* TAC125 to severe nutrient stress. 12 replicate populations were started from single colonies of the wild-type strain and evolved in minimal marine seawater supplemented with 0.1% D-gluconic acid for 200 generations. Serial transfers were performed using a 1:100 dilution factor every two days for 30 days and subsequently every day for 15 days. The experiment was performed using 2 mL of medium with constant shaking at 15 °C.

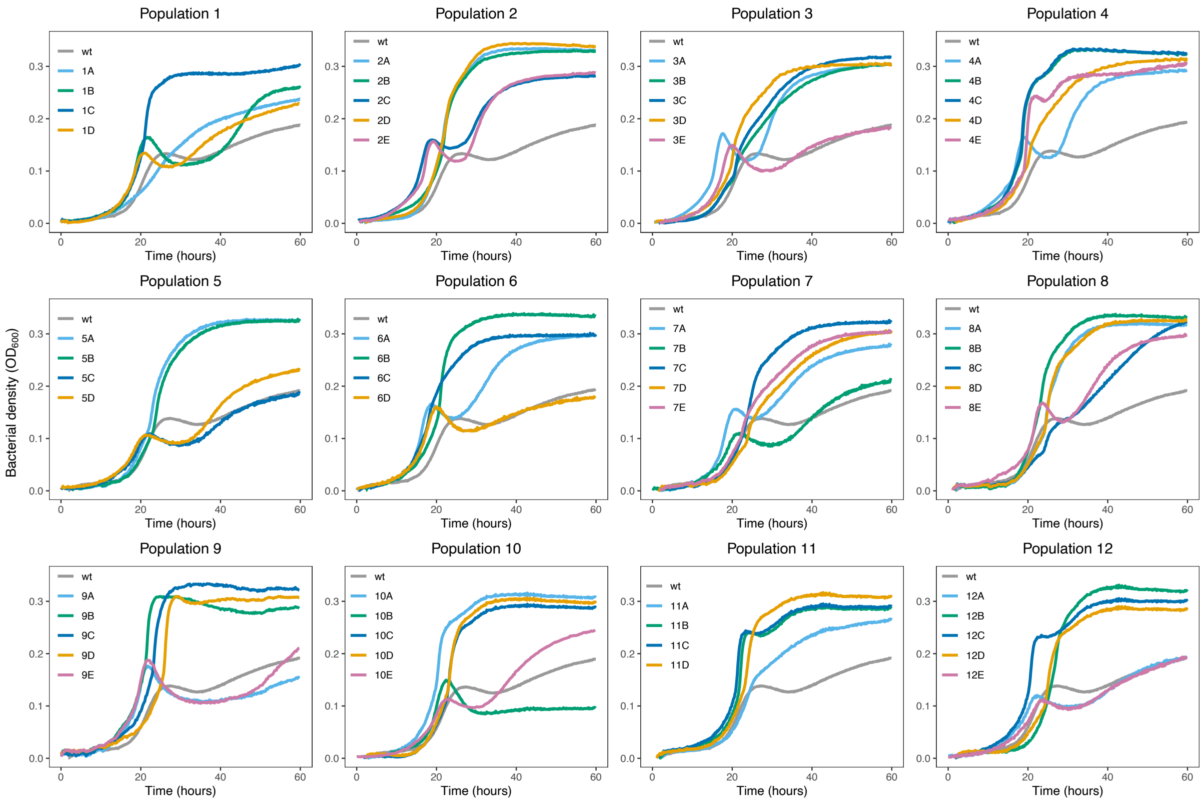


Supplementary Figure 2. Growth curves of the clones selected for whole genome sequencing from each population. Cultures were grown in minimal marine sea water medium supplemented with 0.1% D-gluconic acid at 15 °C. The x-axis indicates time (in hours) and the y-axis bacterial density ($OD_{600}$). The growth curves represent the average bacterial growth observed for three technical replicates. The growth curve of the wild-type clone is coloured in grey. Growth curves were done for 60 hours, but during the evolution experiment the cultures were transferred every 48 hours for the first 15 transfers, and every 24 hours for the last 15 transfers.

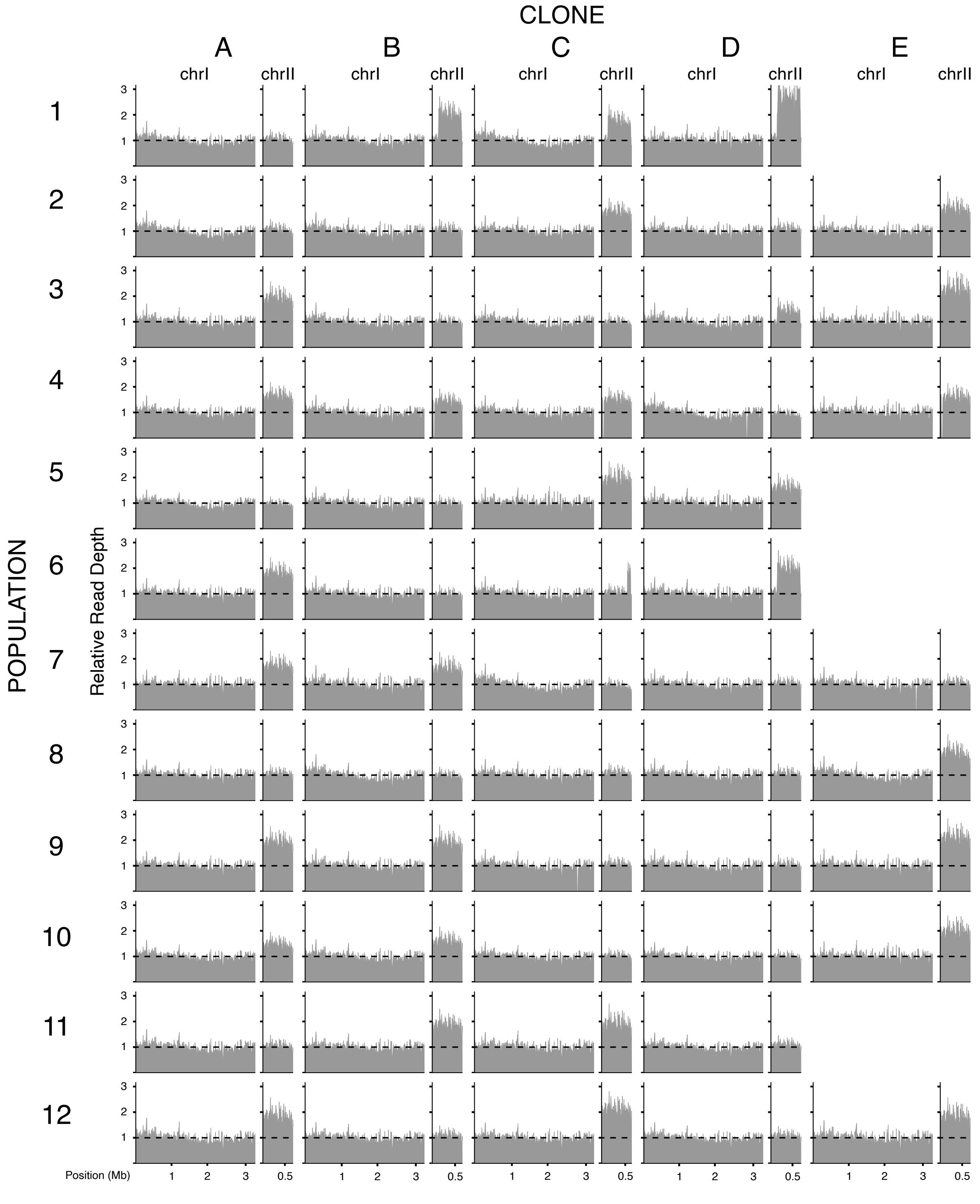


Supplementary Figure 3. Relative read depth in chrI and chrII across sequenced clones. Mean read coverage normalized to chrI for all sequenced clones. Read depth calculated for 5 kb windows. A dashed black line indicates the expected relative coverage when no copy number increments are present. No large deletions of chrI that could affect the chrII/chrI ratio are observed.

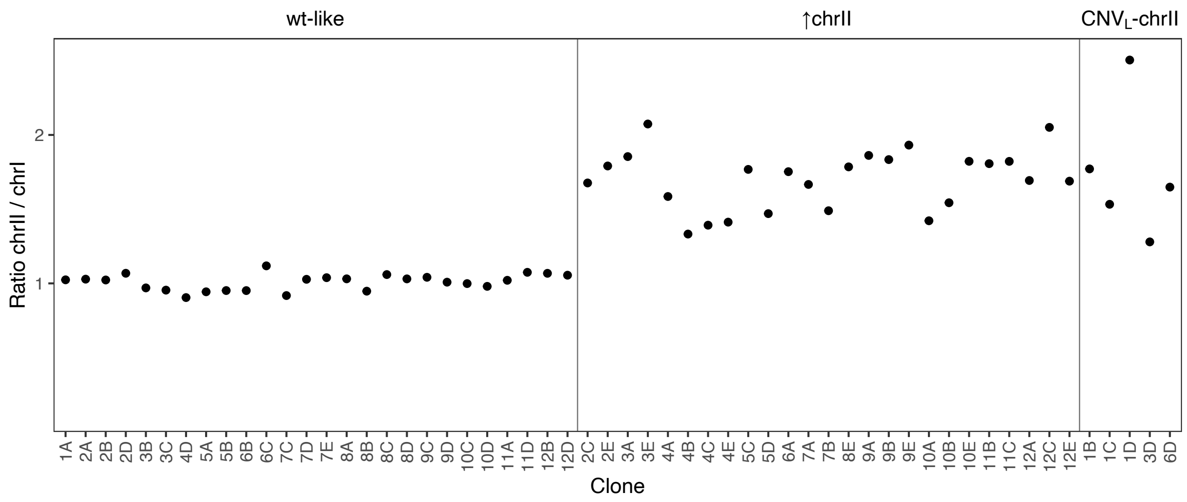


Supplementary Figure 4. Copy number estimation of chrII. The chrII/chrI ratio is computed dividing the mean read coverage of chrII by the mean read coverage of chrI, and then normalized by 0.72 (see methods). Increments in the chrII copy number are assumed when clones have ratios greater than 1.25. Clones with a $CNV_L$-chrII are also shown separately on the right side panel.

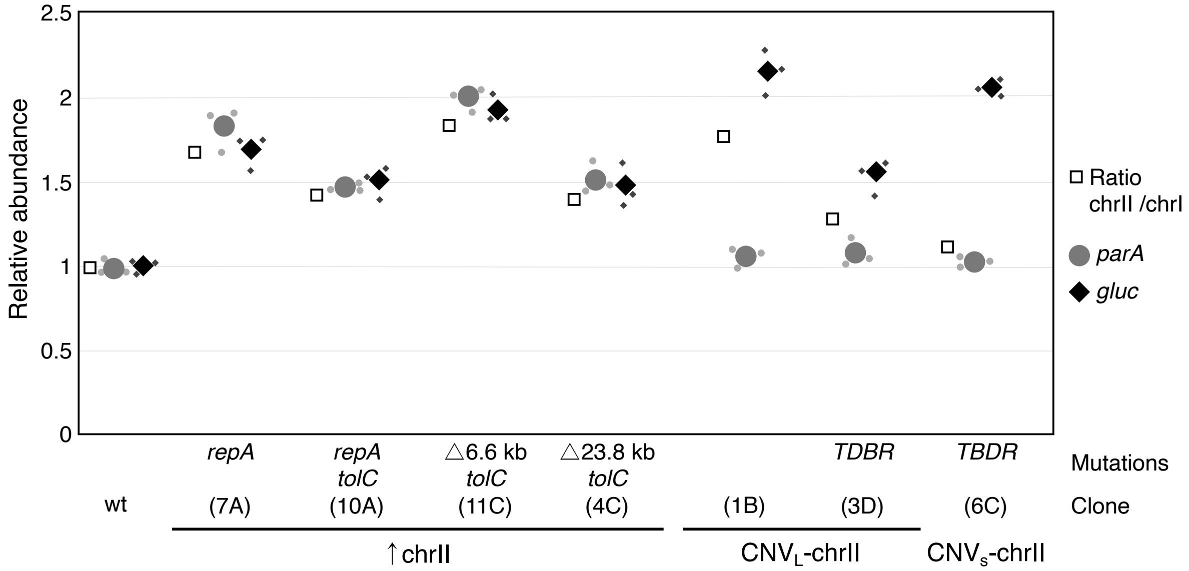


Supplementary Figure 5. Estimation of the copy number of chrII using the chrII/chrI ratio and quantitative PCR. We used qPCR to estimate the relative abundance of two genes (*parA* and *gluc*) located at distant positions in chrII to infer the copy number of chrII across several clones and also to verifty the presence of CNVs. The relative abundance is calculated relative to two reference genes (*ropB* and *gyrA*) located in chrI. Grey circles and black diamonds show the mean relative abundance obtained from three biological replicates (small shapes) for *parA* and *gluc* gene respectively. The square indicates the ratio computed using the number of reads.

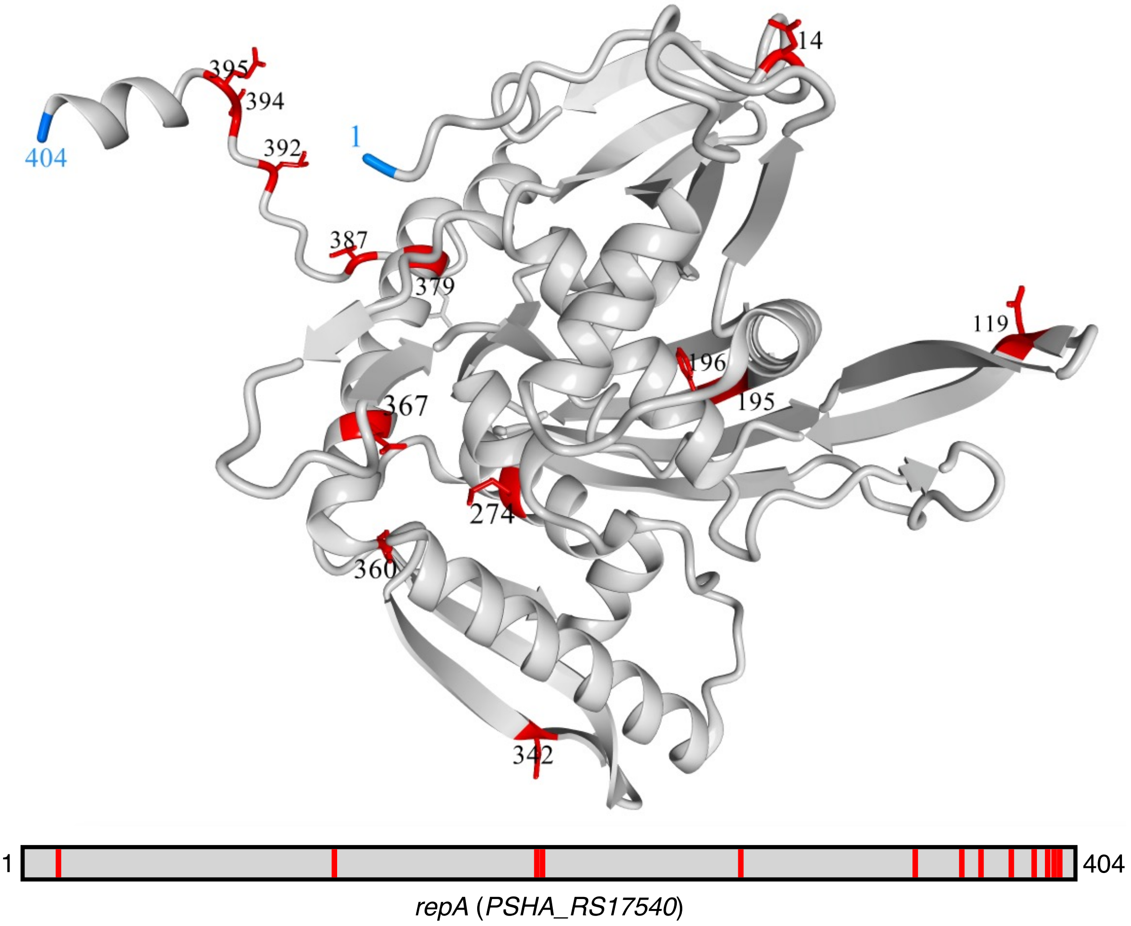


Supplementary Figure 6. Localization of mutations in the tridimensional structure of the RepA protein. Positions affected by mutations are marked in red. The start and end positions of the protein are depicted in blue. We obtained the sequence and the predicted structure of the RepA protein (PSHA_RS17540 gene) of *P. haloplanktis* TAC125 from the AlphaFold Protein Structure Database repository (Q3ICK7)[93]. We visualized and marked the positions of the mutations using Yasara[94].

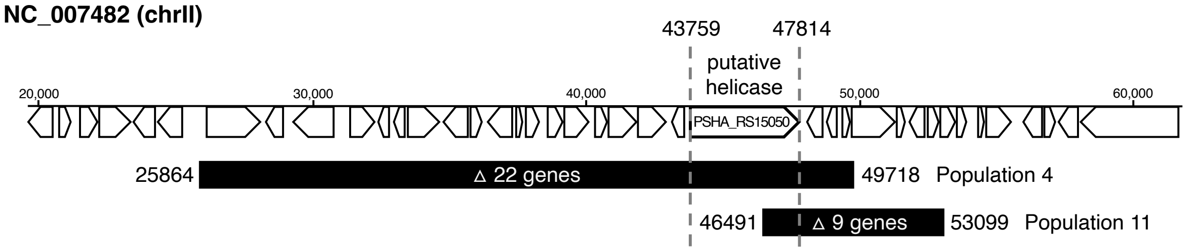


Supplementary Figure 7. Deletions affecting a putative helicase gene are associated to an increase in the copy number of the entire chrII. Clones isolated from populations 4 and 11 carry mutations that involve the complete deletion or truncation of the PSHA_RS15050 gene. The scheme depicts the genes located in the region of chrII hit by the two types of deletions (black rectangles) and the genomic coordinates involved.

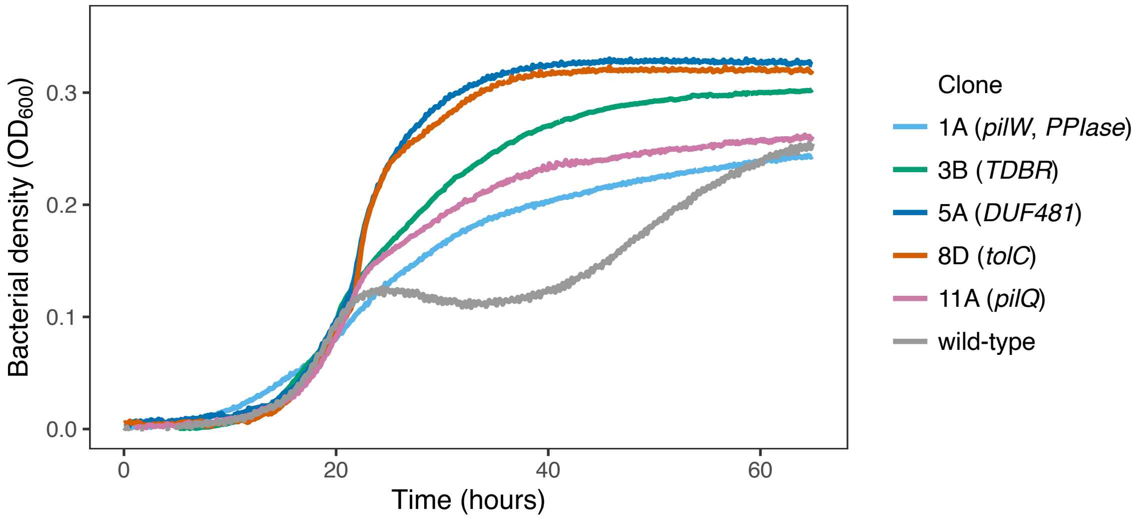


Supplementary Figure 8. Clones with mutations in genes coding for outer membrane proteins and components of the type IV pili show monophasic growth curves. Growth curves performed for clones with single mutations in *tolC*, *TBDR*, *DUF481*, and *PilQ,* and a clone with a mutation in *PilW* together with a mutation in a peptidylprolyl isomerase (PPIase) gene. Cultures were prepared with a starting $OD_{600}$ of 0.005 using MMSW supplemented with D-gluconic acid (0.1%) and bacterial growth (y-axis) was periodically measured in an Infinite 200 Pro (Tecan) microplate reader. The growth curves represent the average bacterial growth observed for three technical replicates.

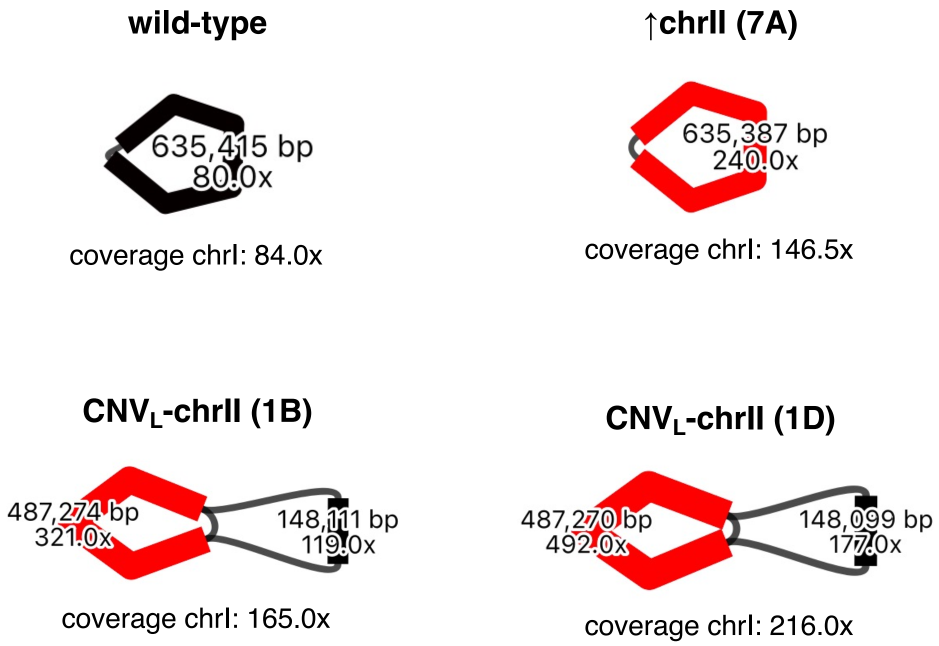


Supplementary Figure 9. Assembly of the chrII in selected clones and the wild-type clone using Oxford Nanopore Sequencing (ONT). Long-reads were assembled into contigs and evaluated to confirm the existence of $CNV_L$-chrII as a tandem duplication inserted into the chrII. We used Bandage for the visualization. Coverage and length for each contig are indicated. Black thin lines represent the regions where contigs can be connected or circularized. Contigs in red indicate a higher relative depth.

**CNV$_L$-chrII**

End (619,758) AGTTATTACAGCCCATAAATTACGTTATGTTAAAT
Start (132,527) TGCACACATATTTTATAAATAAACATTATGTTAAA

CNV$_L$-chrII AGTTATTACAGCCCATAAATAAACATTATGTTAAA

**CNV$_S$-chrII**

End (609,287) TCCTACCATTGAATCTGCTGGTGGCTCAGTAAGGT
Start (539,869) AGATATTACTGAGCCTGCTGGGTATCAAACAATGC

CNV$_S$-chrII TCCTACCATTGAATCTGCTGGGTATCAAACAATGC

Supplementary Figure 10. Sequence context of CNVs in chrII. New junctions created for the CNV$_L$-chrII in clones 1B, 1C, 1D, 3D, and 6D, and for the CNV$_S$-chrII in clone 6C. Orange characters indicate the microhomology shared between the sequences flanking the start and end regions of the CNVs. The sequence flanking the start region is shown in blue while the one flanking the end is coloured in green.

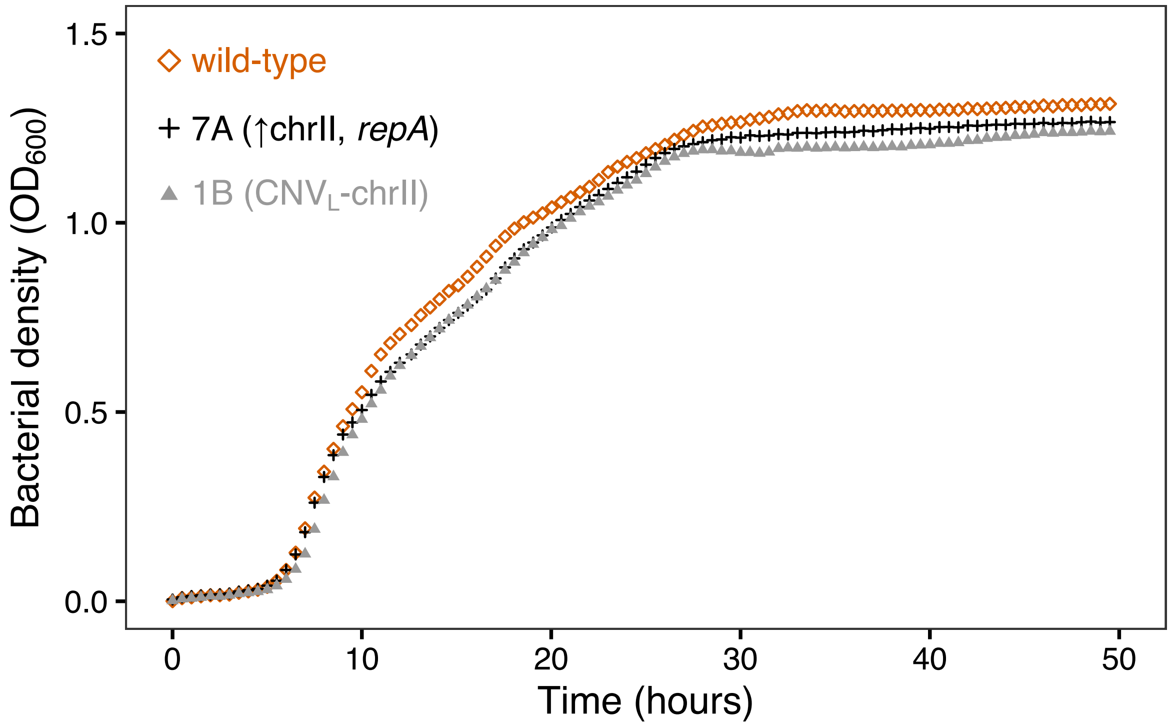


Supplementary Figure 11. Growth comparison in rich medium of clones with changes in chrII copy number and the wild-type clone. The growth of a clone with increased copy number of the entire chrII (7A) and one with the CNV$_L$-chrII (1B) is marginally worst in marine broth 2216 compared to the wild-type clone. We used an Infinite M Nano (Tecan) microplate reader to periodically measure the bacterial growth (y-axis) at 600 nm ($OD_{600}$). Growth curves represent the mean of three biological replicates.

Table S1. Growth type for the ten clones isolated from each of the 12 endpoint populations of the evolution experiment. 10 clones per population were grown in MMSW supplemented with 0.1% D-gluconic acid and classified depending on the type of growth curve observed.

| Population | Biphasic | Monophasic |
|---|---|---|
| 1 | 5 | 5 |
| 2 | 3 | 7 |
| 3 | 4 | 6 |
| 4 | 2 | 8 |
| 5 | 6 | 4 |
| 6 | 5 | 5 |
| 7 | 4 | 6 |
| 8 | 2 | 8 |
| 9 | 3 | 7 |
| 10 | 4 | 6 |
| 11 | 0 | 10 |
| 12 | 4 | 6 |
| Total | 42 | 78 |

Table S2. List of mutations identified in clones isolated from endpoint populations. We mapped the sequenced genomes to the *P. haloplanktis* TAC125 wild-type reference genome to identify mutations gained during the evolution experiment.

| Replicon | Total |
|---|---|
| Chromosome I | 45 |
| Chromosome II | 32 |
| pMEGA | 0 |
| pMtBL | 0 |
| Total | 77 |

| Type of mutation | Effect | Total | Total |
|---|---|---|---|
| Single-base substitutions | | 33 | |
| | Intergenic | | 0 |
| | Missense variant | | 32 |
| | Nonsense variant | | 1 |
| | Synonymous variant | | 0 |
| Indels < 35 nt | | | |
| Insertion | | 19 | |
| | In-frame insertion | | 18 |
| | Frameshifting variant | | 0 |
| | Intergenic region | | 1 |
| Deletion | | 9 | |
| | In-frame deletion | | 0 |
| | Frameshifting variant | | 9 |
| | Intergenic region | | 0 |
| Deletions > 35 nt | | 10 | |
| | In-frame deletion | | 2 |
| | Several genes | | 8 |
| Duplications | | 6 | |
| Total | | 77 | |

Table S3. Loss of $CNV_L$-chrII in colonies obtained from clone 1B. The presence of $CNV_L$-chrII was evaluated by PCR amplification of the new junction generated after the CNV insertion. Clone 1B was grown on agar plates containing MMSW and 0.1% D-gluconic acid and 31 individual colonies were isolated to assess the presence of $CNV_L$-chrII. Three biological replicates were performed.

| Sample | Loss of $CNV_L$-chrII |
|---|---|
| Replicate 1 | 21.05 % |
| Replicate 2 | 28.12 % |
| Replicate 3 | 22.58 % |

Table S4. List of primers used in this study.

| Primer name | Sequence | Purpose |
|---|---|---|
| iC2_4 | ACAGGGCATCTGGATCTTATACT | PCR. Forward primer to amplify the new junction in clones with the $CNV_L$-chrII |
| iC2_6 | AACAGGGAGTAAGCCGGTAA | PCR. Reverse primer to amplify the new junction in clones with the $CNV_L$-chrII |
| rpoB_F | GTGCGTGTAGAACGTGCTGT | qPCR. Forward primer for *rpoB* (PSHA_RS01095) in chrI |
| rpoB_R | AACTGAGACGAGCCGAAGAA | qPCR. Reverse primer for *rpoB* (PSHA_RS01095) in chrI |
| gyrA_F | AAAATGCCTGAAGGACAACG | qPCR. Forward primer for *gyrA* (PSHA_RS06995) in chrI |
| gyrA_R | CGACTCTTCGCTGGGTAGTC | qPCR. Reverse primer for *gyrA* (PSHA_RS06995) in chrI |
| gluc_F | GCGAGAAAAACGCTAACTGG | qPCR. Forward primer for *gluc* (PSHA_RS17145) in chrII |
| gluc_R | ACACTACCACCGCCTTTACG | qPCR. Reverse primer for *gluc* (PSHA_RS17145) in chrII |
| parA_F | GCTGAAGTTGCCGAAAAGTC | qPCR. Forward primer for *parA* (PSHA_RS14855) in chrI |
| parA_R | TCGTGGCGTCTTTTTAAGGT | qPCR. Reverse primer for *parA* (PSHA_RS14855) in chrI |

Table S5. Sequencing output metrics obtained from the Oxford Nanopore Sequencing (ONT). Summary result for four clones: the wild type, one clone with increased copy number of the entire chrII (7A), and two clones with $CNV_L$-chrII (clones 1B, 1D).

| Sample | Total reads | Longest read (bp) | Coverage |
|---|---|---|---|
| Wild type | 72,076 | 77,055 | 90 X |
| Clone 7A | 141,228 | 66,152 | 173 X |
| Clone 1B | 157,361 | 78,914 | 198 X |
| Clone 1D | 207,155 | 67,756 | 266 X |
