## Supplementary_Data1_Vega-Cabrera_etal for "Chromosomal plasticity can drive rapid adaptation in bacteria"

**Data file 1. List of mutations identified in all sequenced clones across populations. Mutations relative to the wild-type *P. haloplanktis* TAC125 reference genome were detected using breseq.**

Clone

| Chromosome | Position | Mutation | Chr | 1A | 1B | 1C | 1D | 2A | 2B | 2C | 2D | 2E | 3A | 3B | 3C | 3D | 3E | 4A | 4B | 4C | 4D | 4E | 5A | 5B | 5C | 5D | 6A | 6B | 6C | 6D | 7A | 7B | 7C | 7D | 7E | 8A | 8B | 8C | 8D | 8E | 9A | 9B | 9C | 9D | 9E | 10A | 10B | 10C | 10D | 10E | 11A | 11B | 11C | 11D | 12A | 12B | 12C | 12D | 12E | Total | Annotation | Gene | Description |
|---|---|---|---|---|---|---|---|---|---|---|---|---|---|---|---|---|---|---|---|---|---|---|---|---|---|---|---|---|---|---|---|---|---|---|---|---|---|---|---|---|---|---|---|---|---|---|---|---|---|---|---|---|---|---|---|---|---|---|---|---|---|---|---|
| NC_007481 | 415,725 | A→T | I ||||||||||||||||||||||||||||||||||||||||||1|||||||||1||||| | 2 | N99Y (AAT→TAT) | *PSHA_RS01970* → | DUF481 domain-containing protein |
| NC_007481 | 415,759 | T→C | I ||||||1||||||||||||||||||||||||||||||||||||||||||||||||||| | 1 | F110S (TTC→TCC) | *PSHA_RS01970* → | DUF481 domain-containing protein |
| NC_007481 | 415,764 | T→G | I ||||||||||||||||||||1|1||||||||||||||||||||||||||||||||||| | 2 | Y112D (TAT→GAT) | *PSHA_RS01970* → | DUF481 domain-containing protein |
| NC_007481 | 415,813 | T→A | I ||||||||||||||||||||||||||||||1|||||||||||||||||||||||||||| | 1 | V128E (GTA→GAA) | *PSHA_RS01970* → | DUF481 domain-containing protein |
| NC_007481 | 1,046,498 | Δ1 bp | I |||||||||||||||||||||||||||||||||1|||||||||||||||||||||||| | 1 | coding (739/1518 nt) | *PSHA_RS04915* → | monovalent cation/H+ antiporter subunit D |
| NC_007481 | 1,055,200 | +GAACTTA | I |||||||||||||||||||||||||||||||||||1|||||||||||||||||||||| | 1 | intergenic (-107/+185) | *PSHA_RS04970* ← / ← *PSHA* | M28 family peptidase/hypothetical protein |
| NC_007481 | 2,025,635 | A→C | I ||||||||||||||||||||||||||||||||||1||||||||||||||||||||||| | 1 | L135* (TTA→TGA) | *uvrY* ← | UvrY/SirA/GacA family response regulator transcription factor |
| NC_007481 | 2,025,712 | Δ1 bp | I |||||||||||||||||||||||||||||||||||||||||||||||||||||1||| | 1 | coding (327/651 nt) | *uvrY* ← | UvrY/SirA/GacA family response regulator transcription factor |
| NC_007481 | 2,025,744 | Δ1 bp | I |||||1|||1|||||||||||||||||||||||||||||||||||||||||||||||| | 2 | coding (295/651 nt) | *uvrY* ← | UvrY/SirA/GacA family response regulator transcription factor |
| NC_007481 | 2,082,079 | T→A | I |1||||||||||||||||||||||||||||||||||||||||||||||||||||||| | 1 | K3M (AAG→ATG) | *PSHA_RS09800* ← | peptidylprolyl isomerase |
| NC_007481 | 2,097,539 | Δ90 bp | I |||||||||||||||||||||||||||||||||||||||||||||||||||||||1| | 1 | coding (1873-1962/2643 nt) | *PSHA_RS09840* ← | TonB-dependent receptor |
| NC_007481 | 2,097,715 | C→A | I ||||||||||||1|1|||||||||||||1|||||||||||||||||||||||||||||| | 3 | G596C (GGT→TGT) | *PSHA_RS09840* ← | TonB-dependent receptor |
| NC_007481 | 2,097,721 | C→A | I |||||||||||1|||||||1||||||||||||||1|1|||||||1|1|||||||||||||||||| | 6 | G594C (GGT→TGT) | *PSHA_RS09840* ← | TonB-dependent receptor |
| NC_007481 | 2,099,004 | Δ45 bp | I |||||||||||||||||||||||||||||||||||1|||||||||||||||||||||| | 1 | coding (453-497/2643 nt) | *PSHA_RS09840* ← | TonB-dependent receptor |
| NC_007481 | 2,209,684 | Δ22,355 bp | I ||||||||||||||||||1||||||||||||||1||||||||1|||||||||||||||||| | 3 | 21 genes | *PSHA_RS17835–PSHA_RS1* | 21 genes |
| NC_007481 | 2,737,148 | +AAGCTC | I |||1||1|||1||||||||1|1||1|||||||||||||1|1||1|||||||1||1|1|||1|1|||1|1|| | 16 | coding (565/1356 nt) | *tolC* → | outer membrane channel protein TolC |
| NC_007481 | 2,737,758 | Δ33 bp | I |||||||||||||||||||||||||1||||||||||||||||||||||||||||||| | 1 | coding (1175-1207/1356 nt) | *tolC* → | outer membrane channel protein TolC |
| NC_007481 | 2,879,188 | G→A | I ||||||||||||||||||||||||||||||||||||||||||||||||1|||||||| | 1 | A623V (GCG→GTG) | *PSHA_RS13365* ← | type IV pilus secretin PilQ family protein |
| NC_007482 | 25,864 | Δ23,854 bp | II ||||||||||||||||1|1||1||||||||||||||||||||||||||||||||||||| | 3 | 22 genes | *PSHA_RS14960–[PSHA_RS* | 22 genes |
| NC_007482 | 46,491 | Δ6,608 bp | II |||||||||||||||||||||||||||||||||||||||||||||||||1|1|||||| | 2 | 9 genes | *[PSHA_RS15050]–[PSHA_R* | *[PSHA_RS15050], PSHA_RS15055, PSHA_RS15060, PSHA_RS15065* |
| NC_007483 | 132,527 | +487231 bp | II ||1|1|1|||||||||1|||||||||||||1|||||||||||||||||||||||||||||| | 5 | 524 genes | *PSHA_RS15400–PSHA_17490* | 524 genes (CNV-L-chrII) |
| NC_007482 | 174,762 | A→C | II |1||||||||||||||||||||||||||||||||||||||||||||||||||||||| | 1 | I142S (ATC→AGC) | *pilW* ← | type IV pilus biogenesis/stability protein PilW |
| NC_007483 | 539,869 | +69409 bp | II ||||||||||||||||||||||||||1|||||||||||||||||||||||||||||| | 1 | 65 genes | *PSHA_RS17140–PSHA_17450* | 65 genes (CNV-S-chrII) |
| NC_007482 | 633,533 | C→T | II |||||||||||||||||||||||1||||||||||||||||||||||||||||||||| | 1 | E395K (GAA→AAA) | *PSHA_RS17540* ← | repA |
| NC_007482 | 633,534 | Δ17 bp | II ||||||||||1||||1|||||||||||||||||||||||||||||||||||||||||| | 2 | coding (1166-1182/1215 nt) | *PSHA_RS17540* ← | repA |
| NC_007482 | 633,541 | Δ4 bp | II |||||||||1|||||||||||||||||||||||||||||||||||||||||||||||| | 1 | coding (1172-1175/1215 nt) | *PSHA_RS17540* ← | repA |
| NC_007482 | 633,556 | Δ2 bp | II |||||||||||||||||||||||||||||||||||||1||||||||||||||||||| | 1 | coding (1159-1160/1215 nt) | *PSHA_RS17540* ← | repA |
| NC_007482 | 633,580 | (CAATACGCTT. | II ||||||||||||||||||||||1|||||||||||||||||1||||||||||||||||||| | 2 | coding (1136/1215 nt) | *PSHA_RS17540* ← | repA |
| NC_007482 | 633,616 | T→A | II |||||||||||||||1|||||||||||||||||||||||1|||||||||||||||||||| | 2 | E367V (GAG→GTG) | *PSHA_RS17540* ← | repA |
| NC_007482 | 633,638 | G→A | II ||||||||||||||||||||||||||||||||||||||||||||||||||||||1|| | 1 | P360S (CCA→TCA) | *PSHA_RS17540* ← | repA |
| NC_007482 | 633,692 | T→A | II ||||||||||||||||||||||||||||||||||||||||||1|||||||||||||| | 1 | N342Y (AAT→TAT) | *PSHA_RS17540* ← | repA |
| NC_007482 | 633,894 | C→T | II ||||||||||||||||||||||||||||||||||||||||||||||||||||1||||1 | 2 | M274I (ATG→ATA) | *PSHA_RS17540* ← | repA |
| NC_007482 | 634,128 | A→T | II ||||||||||||||||||||||||||||||||||||||||||||1|||1||||||||| | 2 | F196L (TTT→TTA) | *PSHA_RS17540* ← | repA |
| NC_007482 | 634,132 | G→A | II |||||||1||||||||||||||||||||||||||||||||||||||||||||||||| | 1 | S195L (TCG→TTG) | *PSHA_RS17540* ← | repA |
| NC_007482 | 634,361 | C→A | II ||||||||||||||||||||||||||||1|1|||||||||||||||1||||||||||||| | 3 | D119Y (GAC→TAC) | *PSHA_RS17540* ← | repA |
| NC_007482 | 634,674 | A→T | II ||||||||||||||||||||||||1|||||||||||||||||||||||||||||||| | 1 | D14E (GAT→GAA) | *PSHA_RS17540* ← | repA |
| Total | | | | 2 | 1 | 2 | 1 | 2 | 1 | 1 | 2 | 1 | 1 | 1 | 1 | 2 | 1 | 1 | 2 | 2 | 2 | 2 | 1 | 1 | 1 | 1 | 1 | 1 | 2 | 1 | 1 | 1 | 1 | 1 | 2 | 2 | 2 | 2 | 1 | 1 | 1 | 2 | 2 | 1 | 1 | 2 | 1 | 1 | 1 | 1 | 1 | 2 | 2 | 1 | 1 | 2 | 2 | 1 | 1 | 77 | | | |
| | | | | m | b | m | b | m | m | b | m | b | b | m | m | m | b | b | m | m | m | m | m | m | b | b | b | m | m | b | b | b | m | m | m | m | m | m | m | b | b | m | m | m | b | m | b | m | m | b | m | m | m | m | b | m | m | m | b | | Type of Growth Curve. Blue: monophasic, orange: biphasic | | |
